# Long-range electrostatic forces govern how proteins fold on the ribosome

**DOI:** 10.1101/2025.02.10.637539

**Authors:** Julian O. Streit, Sammy H.S. Chan, Charles Burridge, Joel Wallace, Christopher A. Waudby, Lisa D. Cabrita, John Christodoulou

**Affiliations:** Institute of Structural and Molecular Biology, University College London, London, UK; Department of Biological Sciences, Birkbeck College, London, UK

## Abstract

Protein biosynthesis and folding are tightly intertwined processes regulated by the ribosome and auxiliary factors. Nascent proteins can begin to fold on their parent ribosome but formation of the native state is often inhibited well beyond the emergence of the necessary residues from the exit tunnel. The dominant forces driving this apparent destabilisation have remained unclear. We investigate this here, combining NMR experiments and atomistic simulations of a folded nascent chain on and off the ribosome. While its native structure and internal dynamics remain unaltered, co-translational folding is disfavoured due to intermolecular electrostatic repulsion between the negatively charged ribosome surface and nascent protein. Partially folded intermediates are less destabilised, resulting in their high populations. Specifically, we show that the polypeptide’s net charge is the dominant factor determining nascent folding thermodynamics, with smaller contributions from charge distribution. Consequently, positively charged proteins can fold on the ribosome without populating stable intermediates. These findings reconcile conflicting observations of previous studies and establish the general physical principles underpinning *de novo* protein folding.

## Introduction

Protein synthesis, folding and assembly are tightly coupled and can occur co-translationally on the ribosome^1–5^. In contrast to decades of research on elucidating the principles of protein folding using isolated proteins^6^, the general physical chemistry principles that underpin co-translational folding (coTF) are only beginning to emerge^7^. Nascent chains (NCs) can fold via different pathways compared to isolated polypeptides, with many proteins able to adopt a functionally active native state only when folded co-translationally but not post-translationally^1,8–11^. Conflicting observations have, however, been made on how and to what extent the ribosome modulates folding. While the ribosome appears to suppress stable coTF intermediates in some cases^9,10,12^, intermediates are highly populated for many other proteins with more favourable folding free energies (ΔG_I-U_) than for isolated proteins by up to ∼5 kcal mol^-1^ (refs.^7,11,13–17^). Stable coTF intermediates have been rationalised, in part, by the entropic destabilisation of unfolded NCs near the ribosome surface – an effect caused by its structural expansion and increased solvation^7^ and with only a minor role for ribosome-NC interactions^7,18^. This entropic advantage to form structure is, however, also re-balanced by a less favourable enthalpy of protein folding which is due to the destabilisation of natively folded states^7^. Indeed, net folding free energies (ΔG_N-U_=ΔG_f(N)_) for the native state have been well documented to be higher on the ribosome (ΔΔG_f(N),RNC-iso_ > 0) for several proteins, effectively delaying complete structure formation in favour of unfolded^9,12^ and partially folded conformations^7,13,18–20^. Reconciling these observations by understanding their origins would provide a significant step towards establishing a predictable framework to study coTF.

Multiple theories and models have been proposed to account for the modulated stability of folded NCs on the ribosome, including the stabilisation of the unfolded state through ribosome interactions^9,18^. However, stabilising contributions of direct ribosome-NC interactions^21^ cannot fully account for coTF thermodynamics^7^. Instead, computational studies have predicted that the hydrophobic effect may be reduced near the ribosome surface due to water ordering^22^, and that folded NCs may experience changes in their internal dynamics and conformational entropy near the ribosome^23^. Experimental studies, on the other hand, have indicated that entropic effects of disordered neighbouring domains may (de)stabilise folded domains^17,24^ and that the ribosome may perturb the native structure of the NC^7^. Moreover, several studies have reported electrostatic effects to influence coTF onset, kinetics, and thermodynamics^9,13,18–20,25,26^. It therefore remains unclear which (de)stabilisation mechanisms contribute most significantly to coTF.

Here, we present structures and thermodynamics of a folded ribosome-bound nascent chain complex (RNC) by combining nuclear magnetic resonance (NMR) experiments and all-atom molecular dynamics (MD) simulations. The integrated approach developed permits a systematic assessment of the range of factors modulating the thermodynamic stabilities of structured NCs. We find that long-range electrostatic effects and NC net charge are the dominant determinants of the thermodynamic stabilities of structured NC states. These effects are strongly destabilising for native states of negatively charged proteins, even in the absence of significant ribosome-NC interactions. Our physical model extends to other NC conformational states: partially structured intermediates are less destabilised and thermodynamically favoured, reconciling their common observation, while the structural expansion and entropic destabilisation of unfolded, negatively charged NCs^7^ is rationalised. Conversely, our model identifies the folding pathways of net positively charged proteins to disfavour intermediate states as their unfolded and native states are not destabilised and the dominant species observed co-translationally. Our work therefore reconciles the common but net charge-dependent observation of stable coTF intermediates on the ribosome that underpin *de novo* folding^7,11,13–15^ and misfolding^27–29^ pathways.

## Results

### Folded structure and dynamics are indistinguishable on and off the ribosome

In order to understand the effect of the ribosome on folding energetics, we sought to obtain detailed insights into the structure and dynamics of a folded nascent chain using the well-characterised immunoglobulin-like domain FLN5^7,13,18,30–33^ as the exemplar. We studied its coTF as a series of biosynthetic snapshots using FLN5 with *L* number of linking residues (FLN5+L), comprised of the subsequent domain and a stalling motif^13,34^ (**Fig. 1a**). We examined FLN5 on the ribosome at long linker lengths such that the FLN5 domain is fully emerged from the exit tunnel^35^. The NMR spectra of uniform ^2^H, selectively ^1^H^13^C-methyl L-Ile/L-Leu/L-Val-labelled native FLN5 on and off the ribosome are highly similar with only very small chemical shift perturbations of 0.06 ppm at most (**Fig. 1b**)^31^. Similarly, ^19^F NMR spectra of four FLN5 variants labelled with 4-trifluoromethyl-L-phenylalanine (tfmF) at two solvent-exposed (655tfmF and 726tfmF) and two buried (715tfmF and 727tfmF) labelling sites also show only very modest chemical shift changes of 0.01-0.08ppm for the native state (**Fig. 1c**)^13,36^, alongside broad resonances corresponding to intermediate states^7,13^. To obtain further insights into protein dynamics, we generated structural ensembles of folded FLN5 both in isolation and translation-stalled on the ribosome using extensive, all-atom MD simulations in explicit solvent validated by NMR data (see Methods & Supplementary text, **Figs. S1-S3**).

**Figure 1.**
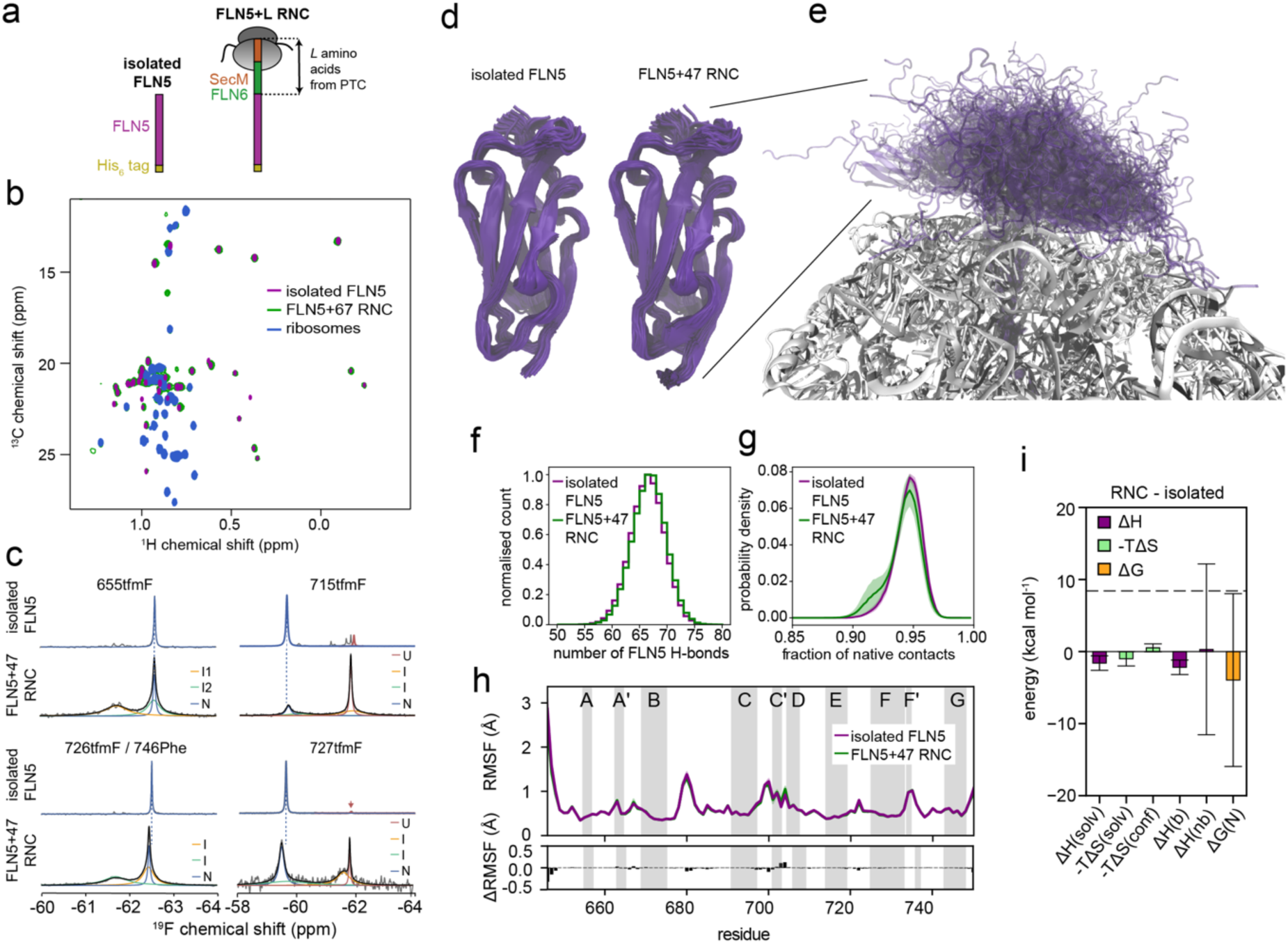
The structure and dynamics of folded FLN5 is not affected by the ribosome. **a.** Schematic of FLN5 constructs in isolation and on the ribosome (FLN5+*L*). **b.** ^1^H,^13^C methyl TROSY NMR spectra of isolated FLN5, FLN5+67 and empty ribosomes recorded at 298 K and 900 MHz^31^ c. ^19^F NMR spectra of FLN5 labelled at different positions with 4-trifluoromethyl-L-phenylalanine (tfmF) recorded at 298K and 500 MHz^13,36^. Observed, fitted and total fitted spectra shown in grey, colour and black, respectively. **d.** Representative structural ensembles of FLN5 in isolation and on the ribosome (FLN5+47) obtained from MD simulations (DES-Amber 3.20). **e.** Structural ensemble of FLN5+47 on the ribosome showing orientational heterogeneity and dynamics. **f.** Histogram of the number of hydrogen bonds (H-bonds) within FLN5 (residues 646-749) calculated from the MD ensembles. **g.** Probability distribution of the fraction of native heavy atom contacts within FLN5 (mean ± s.e.m. from eight independent simulations of 2 μs). **h.** Backbone dynamics quantified by the root mean squared fluctuations (RMSF) of the Cα atoms (mean ± s.e.m. from eight independent simulations of 2 μs). The bottom panel shows the difference (RNC-isolated). **i.** Bar plot summarising predicted energetic changes of folded FLN5 (RNC-isolated protein) from solvation (solv), conformational entropy (conf), bonded (b), and nonbonded (nb) potential energy (see Methods). The dashed, horizontal line represents the lower bound estimate of the native state destabilisation previously determined^7^. All values represent the mean ± s.e.m. from eight independent simulations of 2 μs. The s.e.m. of the conformational entropy term was calculated using a blocking analysis (see Methods).

In line with the experimental chemical shifts, the MD structures of FLN5 in isolation and on the ribosome (FLN5+47) are observed to be highly similar with an RMSD of 1.5 and 1.6 Å relative to the crystal structure, respectively (**Figs. 1d, S2a-b**). The average RMSD of the RNC compared to isolated protein snapshots is 2.0 Å, indistinguishable from the average pairwise RMSD of each ensemble (**Table S1**). On the ribosome, FLN5 samples a highly orientationally heterogeneous and dynamic conformational space with only transient (up to 20%) ribosome interactions (**Fig. 1e, S2e**).

### Energetic origins of nascent chain destabilisation

We considered the hydrogen bonding network within our structural ensembles as this force is the major stabiliser of structure^37^, and we found that the average number of hydrogen bonds is statistically indistinguishable on and off the ribosome (66.2 ± 0.2 and 65.9 ± 0.2 hydrogen bonds, respectively, **Fig. 1f**). Likewise, the fraction of native heavy atom contacts is very similar on and off the ribosome (94.2 ± 0.3 versus 94.6 ± 0.1 %, respectively, **Fig. 1g**). Indeed, an analysis of backbone dynamics showed that the protein behaves identically on and off the ribosome (**Figs. 1h, S2b**). The small chemical shift perturbations observed by NMR are therefore likely due to the difference in environment rather than structure; notably, the largest methyl chemical shift differences are found at the C-terminal hemisphere in closest proximity to the ribosome surface^7,31^. Together, the NMR and MD results strongly suggest that the ribosome does not destabilise folded states by altering the protein structure.

Our previous work on FLN5-RNCs showed a destabilisation of the folded state on versus off the ribosome^7^. This *tethering* free energy difference, ΔG_t,RNC-iso_ (i.e., the free energy difference between the native state on and off the ribosome, G_N,RNC_ – G_N,iso_) was found to be +8.4 kcal mol^-1^ for FLN5+42 (lower bound)^7^. As previously undertaken for the disordered state of FLN5 on the ribosome^7^, here we used the structural ensembles of the folded FLN5+47 RNC described above to consider the energetics to identify the origin(s) that account for this large ΔG_t,RNC-iso._ We predicted the entropic and enthalpic changes FLN5 experiences on the ribosome relative to in isolation (see Methods). On the ribosome, these analyses indicated that transient ribosome interactions slightly reduce FLN5 solvation (**Fig. S4a-c**), resulting in only a mild stabilisation (−2.6 ± 1.4 kcal mol^-1^, **Fig. 1i**). Also, in agreement with the highly similar backbone dynamics observed (**Fig. 1h**), no significant changes in protein conformational entropy were predicted (**Figs. 1i, S4d**). To evaluate the enthalpy of FLN5, we calculated the difference in bonded and nonbonded (van der Waals and electrostatic atomic interactions) potential energy of FLN5 domain atoms (residues 646-749, see Methods). The bonded potential energy shows a mild stabilisation of FLN5 on the ribosome (−2.2 ± 1.0 kcal mol^-1^), whereas the calculated nonbonded potential energy showed a high statistical error of ∼12 kcal mol^-1^ (**Fig. 1i**). These results were also robust to force field modifications that modulate ribosome-NC interactions (‘LJ09’ simulations of FLN5+47, Supplementary text, **Fig. S4f-j**).

These analyses suggest that structural alterations, FLN5 solvation, and conformational entropy cannot account for the destabilisation on the ribosome. A weaker hydrophobic effect near the charged ribosome surface^22^ is also unlikely to account for the destabilisation for proteins that, like FLN5, fold beyond the vestibule (Supplementary text). The entropy of water is reduced within ∼1nm of the ribosome surface, but the folded domain of FLN5+47 occupies a space on average ∼2-2.5 nm from the surface (**Fig. S5**). At these distances, however, the domain may still experience electrostatic (enthalpic) effects, specifically the nonbonded potential energy as the potential mechanism of destabilisation. While the latter cannot be precisely calculated, the uncertainty found (∼12 kcal mol^-1^) is within range of the experimentally determined destabilisation, ΔG_t,RNC-iso_ (**Fig. 1i**). In other words, it may be unfavourable for the negatively charged FLN5 domain to be close to the negatively charged ribosome surface compared to in bulk solution. Additionally, with the emergence of the disordered N-terminus of the subsequent domain, FLN6, linker residues at the C-terminus may affect the stability of the folded domain. Both C-terminal linker and electrostatic effects were further investigated.

### Disordered linker effects do not reconcile coTF thermodynamics

The extension of the NC domain through its disordered linker residues, comprised of the subsequent FLN6 domain, was investigated for its contributions to the native state stability on the ribosome. In previous studies disordered termini have been shown to decrease protein stability^17,24^, while in some cases stability can also be increased^38,39^. The magnitudes of these effects have, however, been modest with changes of up to 1-2 kcal mol^-1^ in general^24,38,39^. To investigate this for FLN5, we measured the thermodynamic stability of isolated FLN5 extended by two linker lengths (+5 and +15 amino acids) in isolation by ^19^F NMR (**Fig. 2a-b**), the latter analogous to the number of residues outside the exit tunnel in FLN5+47 RNC. These extended FLN5 constructs still showed a two-state folding equilibrium in isolation and only modest changes in folding free energy of −0.57 ± 0.02 and −0.24 ± 0.01 kcal mol^-1^, respectively, relative to wild-type (WT) FLN5 (**Fig. 2c**). Both constructs are thus slightly stabilised, and indeed simulations indicate that the linkers are not conformationally restricted and thus not entropically destabilising (**Fig. 2e**). Instead, factors that result in this mild stabilisation may include, amongst others, the neutralisation of C-terminal charge (**Fig. 2d**) and linker-FLN5 interactions which results in a gain in solvent entropy (**Fig. 2f**). Another example of stabilisation by linking residues is the HRAS G-domain^39^, yet we have recently shown HRAS to also be destabilised when tethered to the ribosome by the same linker^7^. Linker effects are therefore not responsible for the large changes in coTF thermodynamics, which must instead originate directly from the ribosome particle.

**Figure 2.**
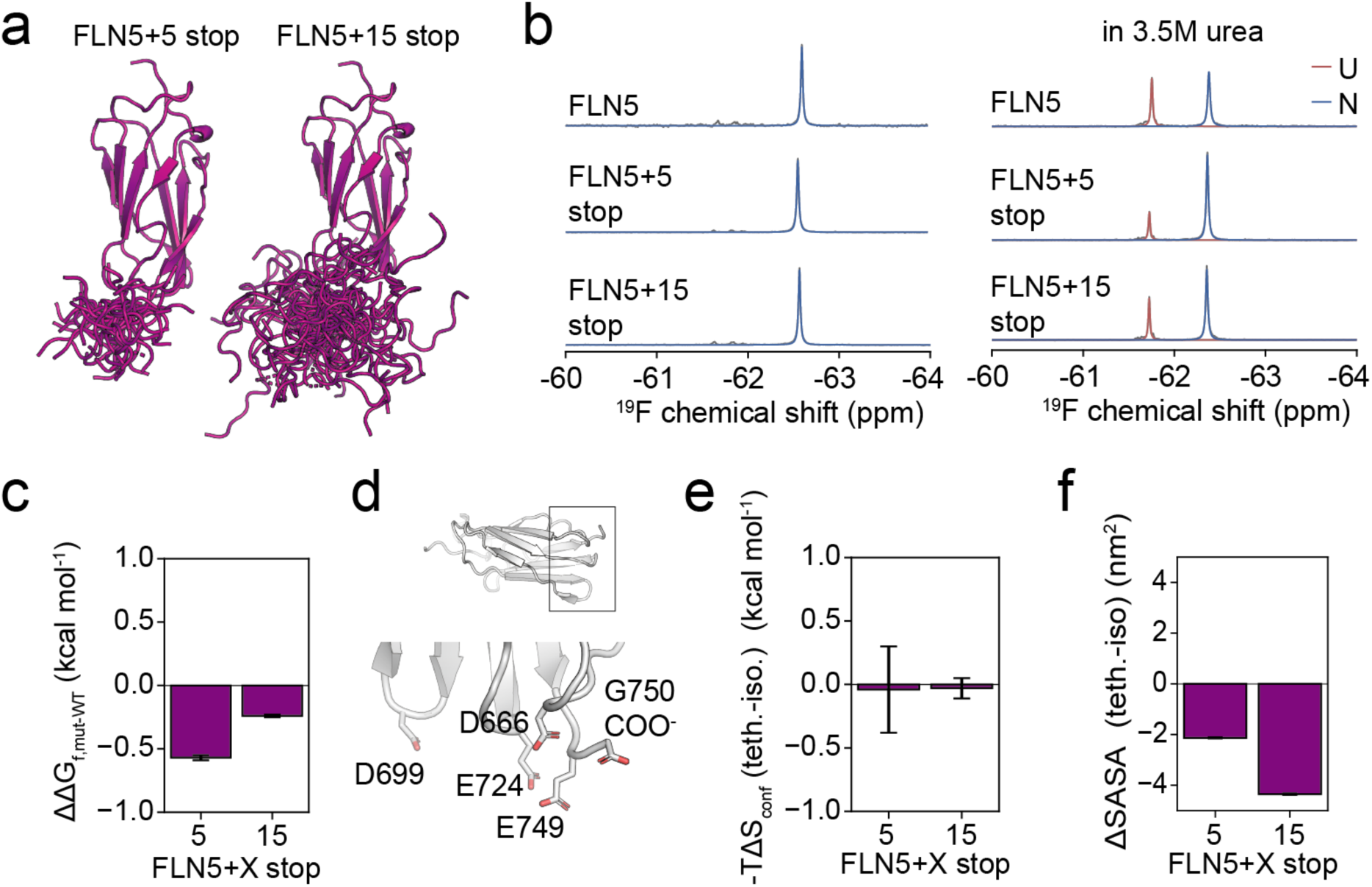
The impact of linker residues on FLN5 native state stability. **a.** Structural models of isolated FLN5 extended by linkers of 5 and 15 residues (FLN5+5 stop and FLN5+15 stop) obtained by coarse-grained MD simulations (see Methods). **b.** ^19^F NMR spectra of isolated FLN5, FLN5+5 stop and FLN5+15 stop (all 655tfmF-labelled) in buffer (left) and in 3.5 M urea (right) recorded at 298 K and 500 MHz. Observed and fitted spectra shown in grey and colour, respectively. **c.** Difference in folding free energy of linker mutants compared to WT FLN5 calculated from the populations of native (N) and unfolded (U) from lineshape analyses of spectra in panel b (mean ± s.e.m.). **d.** Crystal structure of FLN5 (PDB 1QFH) highlighting the abundance of negatively charged group near the C-terminus. **e.** Predicted difference in conformational entropy of the C-terminal linker peptides using coarse-grained MD (mean ± s.e.m., see Methods). **f.** Calculated changes in the solvent-accessible surface area (SASA) of FLN5 and the C-terminal linker relative to their isolated conformations (mean ± s.e.m., see Methods).

### Mutations modulating electrostatics correlate with thermodynamics

We next sought to assess the contributions of ribosome electrostatics, protein net charge, and charge distribution within the NC. Since intermolecular electrostatic effects cannot be precisely quantified from MD simulations (**Fig. 1i**), we investigated these effects experimentally. We reasoned that increasing the net charge of FLN5 would decrease the destabilisation of the native state and designed four charge mutants. Native FLN5 protein has a net charge of −9 at physiological pH, and we designed mutants to each have a net charge of −2 with differing charge distributions throughout the sequence (**Fig. 3a**) and charge patches on the surface of folded FLN5 (**Fig. 3b**). The increased net charge will inevitably also stabilise the unfolded state by promoting ribosome interactions with positively charged unfolded regions^7,18,40^, resulting in a compaction and entropic stabilisation of the unfolded state (**Fig. S6a-c**)^7,9^. However, if the effects of the folded state dominate then the folding free energy will become more favourable and provide direct evidence for native state stabilisation by decreasing the intermolecular electrostatic repulsion from the ribosome surface.

**Figure 3.**
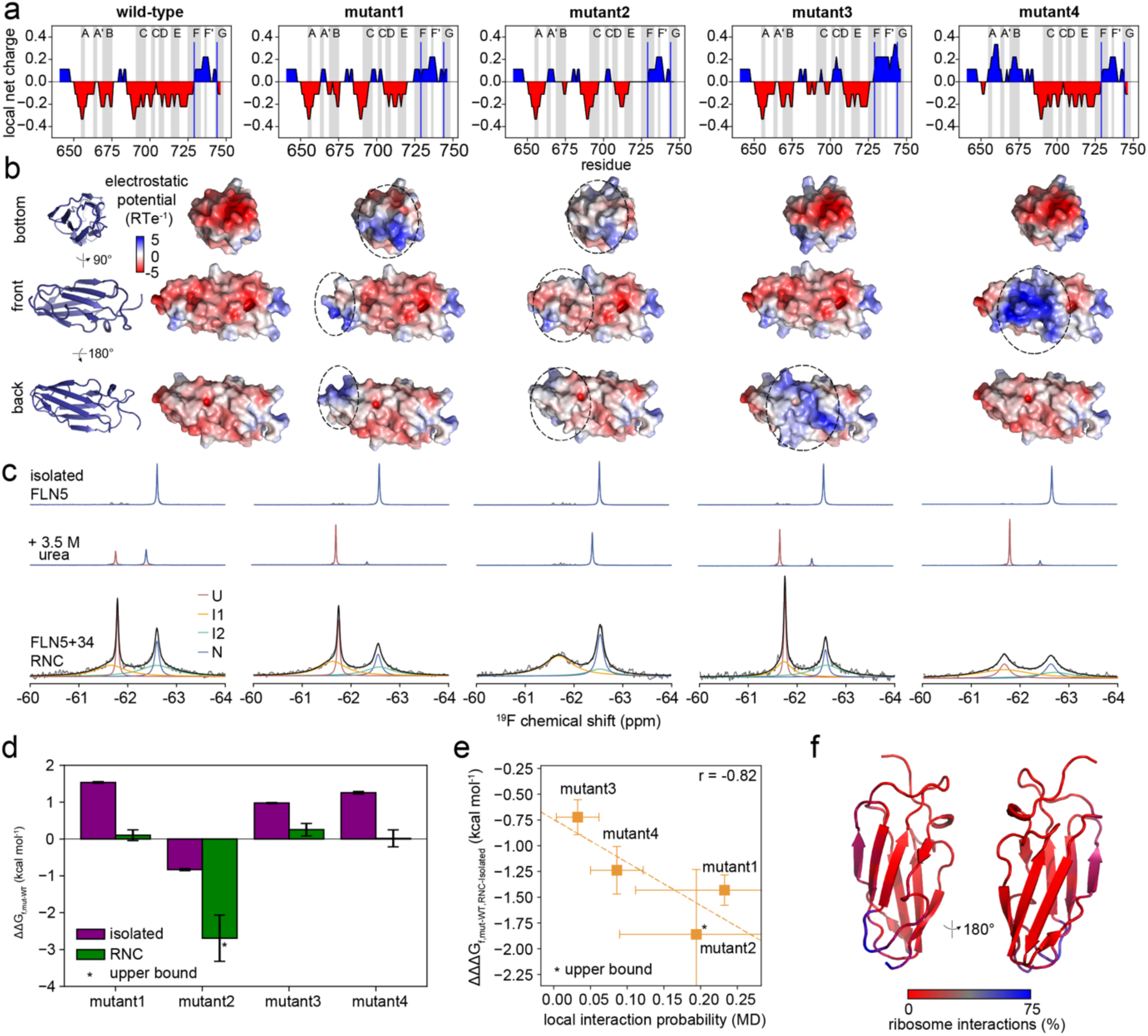
Modulating electrostatic effects with charge mutations. **a.** Local net charge of all mutants along the sequence of FLN5 averaged over a window size of nine residues. β-strand regions are annotated, and blue lines highlight the C-terminal region that preferentially interacts with the ribosome^18^. **b.** Surface electrostatic potentials of all FLN5 variants shown from three viewpoints. Differences relative to WT FLN5 are circled. **c.** ^19^F NMR spectra of FLN5 655tfmF variants in isolation (top and middle, in the presence and absence of urea, respectively) and on the ribosome (FLN5+34, bottom) recorded at 298 K and 500 MHz. Observed, fitted and total fitted spectra shown in grey, colour and black, respectively. **d.** Changes in folding free energies (ΔΔG_f,mut-WT_) measured by ^19^F NMR for all mutants relative to WT FLN5 on and off the ribosome. Errors represent the standard error of the mean obtained from bootstrapping of lineshape fits. The folding free energy of mutant 2 in isolation was calculated from a spectrum recorded in 4.5 M urea to observe the unfolded state (Fig. S7). **e.** Correlation of change in mutant (de)stabilisation (ΔΔΔG_f,mut-WT,RNC-Isolated_) and the average interaction probabilities of residues within 1 nm of a mutant residue (defined by Cα atoms) calculated from MD simulations of wild-type FLN5+34 with the DES-Amber 3.20 force field. Error bars for interaction probabilities represent the standard deviation across all residues contributing to the average. **f.** Average ribosome interaction probability (wild-type FLN5+34 MD, DES-Amber 3.20) mapped on the crystal structure of FLN5 (PDB 1QFH).

We recorded ^19^F NMR experiments^13^ of all mutants and WT FLN5 on and off the ribosome. The FLN5+34 RNC spectra showed four states populated at equilibrium (unfolded, two intermediates, I1 and I2, and the native state) while the isolated proteins in urea only showed two resonances corresponding to the unfolded and native state, consistent with previous work^7,13^, and therefore permitted direct quantification of the folding free energy, ΔG_f_, from the integrals of the unfolded and folded resonances (**Figs. 3c, S7**). Mutants 1 (N667K/D699K/E724K/E749K), 3 (E692K/D704K/N730K/D744K) and 4 (E657K/E659K/E671K/T673K) are destabilised in isolation by +1.0-1.5 kcal mol^-1^, while mutant 2 (D666N/E671Q/D699N/D704N/D720N/E724Q/E749Q) is stabilised by −0.8 kcal mol^-1^ (**Figs. 3c, S7**). However, as expected, on the ribosome all mutants were less destabilised (mutants 1,3 and 4) or more stabilised (mutant 2) compared to in isolation (**Fig. 3d**). While we have previously reported that destabilisation by mutation is generally buffered on the ribosome by up to 70% in magnitude for mutants with the same net charge, the reductions in ΔΔG_f,mut-WT_ are substantially stronger (75-99%). This clearly shows that electrostatic forces exert an effect in addition to mutation buffering, and indeed mutant 2 confirms this. The magnitude of ΔΔG_f,mut-WT_ for the stabilising mutant 2 was increased to at least −2.7 ± 0.6 kcal mol^-1^ on the ribosome (**Fig. 3d**). These results therefore support the physical model that folded FLN5 is destabilised due to its highly negative net charge and an unfavourable intermolecular potential energy term on the ribosome.

In our considerations of the role of charge distribution in the observed stabilities, we intriguingly found that the difference in mutant (de)stabilisation on and off the ribosome (ΔΔΔG_f,mut-WT,RNC-Iso_) correlates with the ribosome interaction probability (calculated form our MD simulations) of the structural regions that were mutated (**Fig. 3e-f, S8**). For example, residues bearing the mutations in mutant 3 interact the least with the ribosome, and the stability of this variant is the least affected by the mutations. The correlation can be rationalised by charged regions with shorter ribosome distances having a greater contribution to the potential energy. This analysis further shows that charge distribution contributes to coTF energetics, which are related to the structural and orientational preferences of the emerging NC.

### Co-translational native state stability is determined by electrostatics

We sought to further deconvolute the long-range electrostatic effects on folded and unfolded conformations. Ionic strength can influence NC dynamics, folding kinetics, and thermodynamics^9,13,18–20^ although it has remained unclear whether these observed effects originate from the unfolded or folded states, or both. The C-terminus of FLN5 binds to the ribosome surface due to a polybasic patch (**Fig. 4a**), stabilising the unfolded state by ∼1 kcal mol^-1^ at FLN5+34^18^ and shifting the equilibrium towards the folded state with increasing ionic strength^13,18^. We therefore sought to examine the folding free energy in response to changing ionic strength in a variant of FLN5 where the unfolded state contributions are negligible. We thus selected the “E6” variant, with six glutamate mutations in the C-terminal binding region (**Fig. 4a-b**), which reduces unfolded state binding^18^. The amount of residual unfolded state binding (∼10%) can only shift folding towards native FLN5 by less than 0.1 kcal mol^-1^ (ΔΔG_f_ = RTln(1-p_B_))^18^.

**Figure 4.**
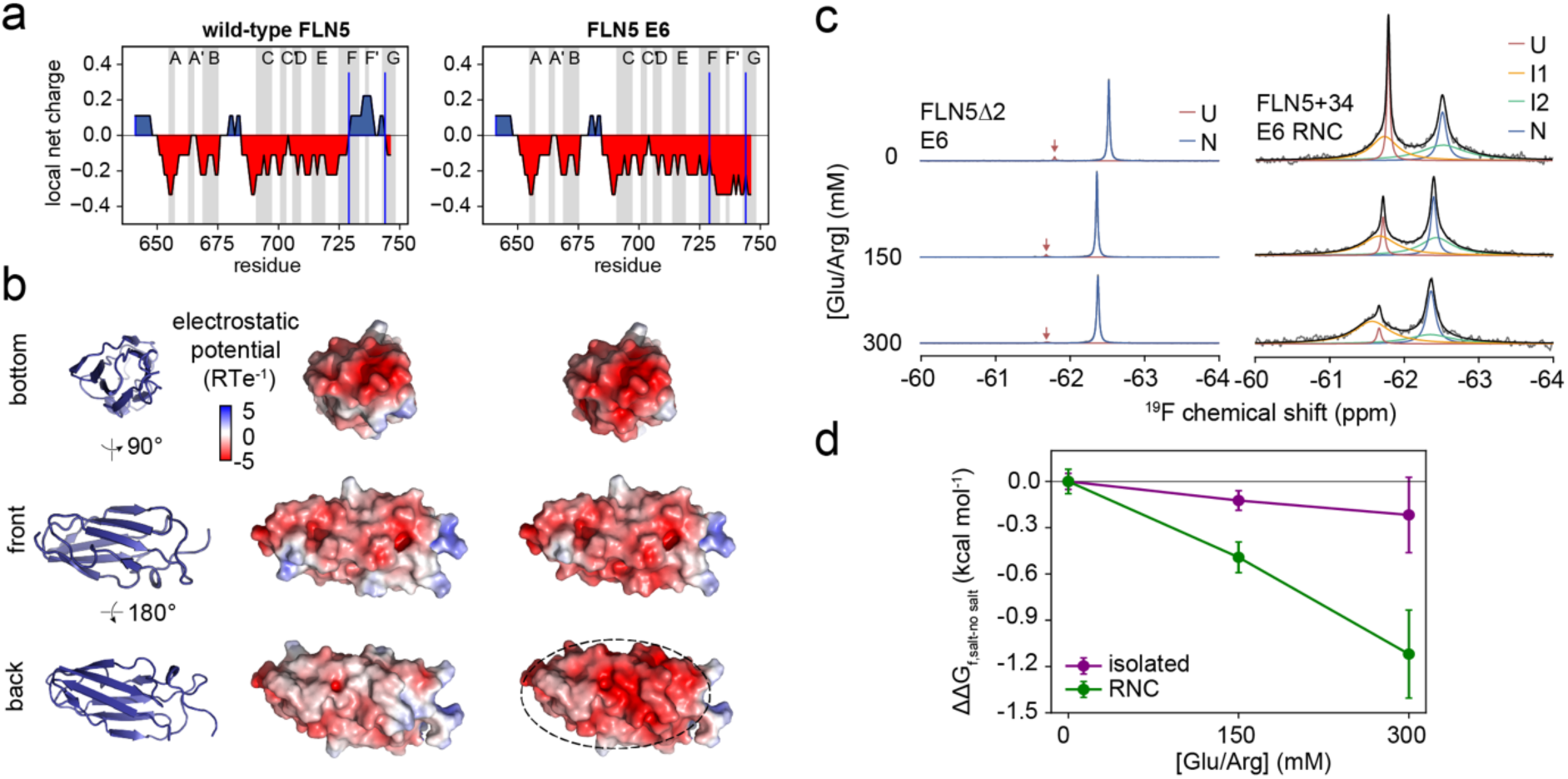
Effect of ionic strength on native state stability. **a.** Local net charge of FLN5 WT and E6 averaged over a window size of nine residues. β-strand regions are annotated, and blue lines highlight the C-terminal region that preferentially interacts with the ribosome^18^. **b.** Surface electrostatic potentials of FLN5 WT and E6 shown from three viewpoints. Differences relative to WT FLN5 are circled. **c.** Salt (L-Glu/L-Arg) titration effects on ^19^F NMR spectra of FLN5 E6 655tfmF in isolation (top, FLN5Δ2^7^) and on the ribosome (FLN5+34, bottom) recorded at 298 K and 500 MHz with increasing ionic strength. Observed, fitted and total fitted spectra shown in grey, colour and black, respectively. **d.** Changes in folding free energies (ΔΔG_f,salt-no salt_) measured by ^19^F NMR as a function of salt concentration. Errors represent the standard error of the mean obtained from bootstrapping of lineshape fits.

We measured the folding free energy of FLN5 E6 in isolation and in FLN5 E6+34 on the ribosome with increasing ionic strength by ^19^F NMR (**Figs. 4c, S9**). In the presence of 300 mM L-Glu/L-Arg, the domain is stabilised by −0.9 ± 0.4 kcal mol^-1^ more on compared to off the ribosome, substantially more than expected from disrupting residual interactions with the unfolded state (< 0.1 kcal mol^-1^, **Fig. 4d**). At higher ionic strength (450 mM L-Glu/L-Arg), folding is even further favoured with no measurable population of unfolded state, although this limits quantification of free energy changes (**Fig. S9**). These data therefore support the electrostatic destabilisation model of predominantly folded, rather than unfolded, FLN5 and show that this destabilisation can be rescued by increasing the ionic strength.

### Electrostatic effects on partially folded intermediates

We next investigated whether the partially folded coTF intermediates of FLN5 experience similar electrostatic effects to the native state. We further analysed the ^19^F NMR data and extracted folding free energies from the mutant (**Fig. 3**) and salt titration experiments (**Fig. 4**). The mutants that increase FLN5 net charge and stabilise the native state also stabilise both coTF intermediates, I1 and I2 (**Fig. 5a**). Likewise, increasing the ionic strength stabilises both I1 and I2 (**Fig. 5b**). These data thus clearly show that partially folded intermediates, like the native state, are destabilised on the ribosome (ΔG_t,I_ > 0) due to unfavourable electrostatic forces. This destabilisation is attenuated by increasing the domain’s net charge and the ionic strength. All FLN5 NC conformations are therefore destabilised when tethered to the ribosome relative to their isolated counterparts. This includes the unfolded state, as we have previously shown it to be entropically destabilised via its structural expansion that results in increased solvation^7^.

**Figure 5.**
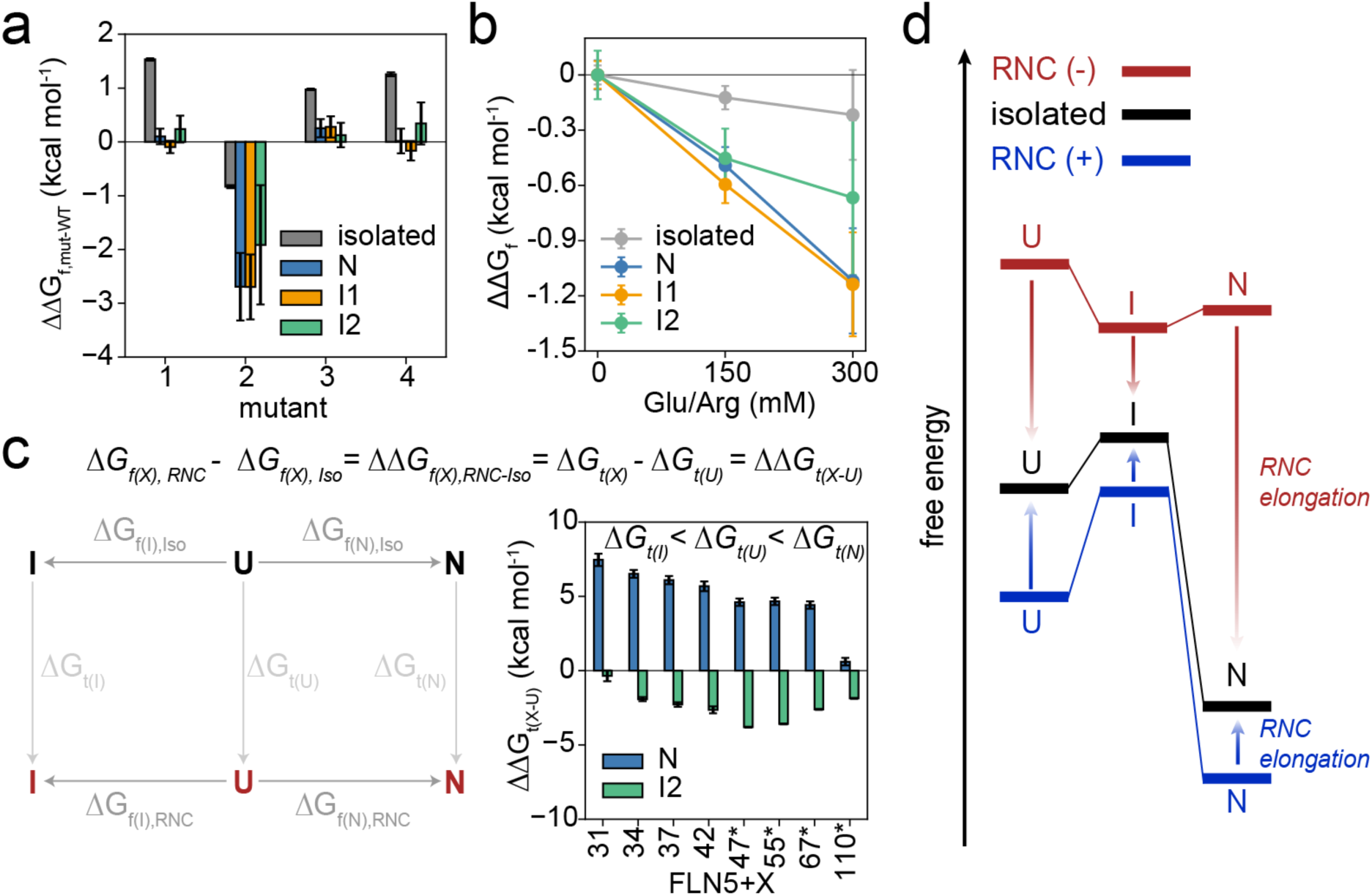
The ribosome distorts the free energy landscape to favour partially folded intermediate states for negatively charged proteins. **a.** Changes in folding free energies (ΔΔG_f,mut-WT_) measured by ^19^F NMR for all mutants and all structured RNC states relative to WT FLN5 on and off the ribosome. Errors represent the standard error of the mean obtained from bootstrapping of lineshape fits. **b.** Changes in folding free energies (ΔΔG_f,salt-no salt_) measured by ^19^F NMR as a function of salt concentration for all structured RNC states. Errors represent the standard error of the mean obtained from bootstrapping of lineshape fits. **c.** (Left) Thermodynamic cycles including folding (ΔG_f_) of intermediate (I) and native (N) states from the unfolded (U) state. Black and red represent isolated protein and RNC, respectively, with the free energy difference due to ribosome tethering (ΔG_t_) connecting these states. (Right) Differences in tethering free energies of structured I and N states relative to the U state (ΔΔG_t(X-U)_) obtained from the thermodynamic cycle and experimentally measured folding free energies^7,13^. We used the I2 coTF intermediate due to structural similarity to an intermediate in isolation^7,13^. **d.** Free energy diagram depicting protein domains in isolation relative to their ribosome-bound forms upon complete emergence from the exit tunnel. For negatively charged RNCs, all conformational states are destabilised on the ribosome relative to their isolated forms (ΔG_t(X)_ > 0). The N state is destabilised the most, followed by the U and I state (panel c). The destabilisation effects decrease with increasing linker length (RNC elongation). Positively charged RNCs are stabilised or unaffected in all NC conformations due to favourable electrostatic interactions (ΔG_t(X)_ ≤ 0, see **Table S2**). At very short linker lengths while the domain is still constrained within the exit tunnel, N and I states are highly destabilised due to steric restrictions regardless of net charge^33,41^. Secondary to net charge, charge distribution modulates the magnitude of these effects experienced by individual conformational states (Fig. 3 and refs.^9,18^) and local nascent peptide segments^20^.

The destabilisation of both the native and intermediate states on the ribosome may appear counterintuitive given that coTF intermediates are highly populated^7,13^. To attempt to reconcile this, we constructed a thermodynamic cycle connecting all protein conformations on and off the ribosome (**Fig. 5c**). From this cycle it becomes apparent that the difference in folding free energy of an intermediate or native state on and off the ribosome (ΔΔG_f(X),RNC-Iso_) is equivalent to the difference in tethering free energy penalty (ΔΔG_t(X-U)_), or in other words the relative destabilisation of ribosome-bound conformational states with respect to their isolated counterparts (**Fig. 5c**). To calculate ΔΔG_t(X-U)_, we therefore used previously measured folding free energies across multiple NC lengths of FLN5 across a wide folding transition^13^. This shows that for all NC lengths where the FLN5 domain has fully emerged from the exit tunnel, the native state is destabilised the most, followed by the unfolded and intermediate states (**Fig. 5c**). Intermediate states are therefore effectively stabilised on the ribosome relative to off the ribosome as they are the least destabilised by ribosome tethering. As expected, the destabilisation of the native state decreases with increasing NC length and its population therefore increases during biosynthesis^13^ (**Fig. 5c**). The intermediate state(s) also become less destabilised up to a linker length of FLN5+47 and then their destabilisation becomes more similar to the unfolded state (**Fig. 5c**), resulting in their populations peaking at FLN5+47 and then decreasing in favour of the native state^13^. Moreover, previous measurements of folding enthalpy and entropy also support that the destabilisation effects of all states decrease with increasing linker length^7^. The ribosome thus modulates the NC conformation-specific destabilisation in a length-dependent manner to steer the folding pathway.

### Protein net charge dictates folding pathways and thermodynamics

To understand whether the electrostatically driven modulation of structure stability on the ribosome is generally applicable to the proteome, we surveyed the literature and compiled a list of proteins for which qualitative or quantitative experimental RNC measurements reporting on protein folding thermodynamics or kinetics are available (**Table S2**). For all systems studied with a negative net charge (ten proteins), the native states were less stable on compared to off the ribosome (**Table S2**). Negatively charged systems which have been studied structurally have also revealed the stable population of coTF intermediates not seen in the isolated protein, including FLN5^7,13^, HemK^14,15^, titin I27^7^, HRAS^7^, firefly luciferase^11^, DHFR^17^, and β-galactosidase^16^. This supports the idea that the energetic effects exerted on negatively charged NCs are generally applicable. Such NCs have an entropically destabilised unfolded state^7^ and the native state is enthalpically destabilised (this work and ref.^7^), which together results in folding intermediates that are more stable on compared to off the ribosome.

The native states of proteins with a highly positive net charge, namely T4 lysozyme^9^ (+8) and DNA polymerase IV (DinB, +9)^20^, are not destabilised and their coTF intermediates are lowly populated (**Table S2**). In line with this, studies comparing different proteins or charge variants have found that increasing net charge resulted in more folding and earlier coTF onset (four proteins)^19,26^ (**Table S2**). These studies have reported that the effects on stability are reduced by increasing NC linker lengths (i.e., distance from the ribosome) and ionic strength^9,19,20^, in agreement with our findings (**Figs. 3-4** and refs.^7,13^). In addition to the dominant effects of net charge, local stabilities of NC segments have been observed to correlate with local charge^20^, with positively charged regions unaffected or stabilised by the ribosome (CheY and DinB, **Table S2**). This is consistent with the modulation of NC stability by charge distribution observed for FLN5 (**Fig. 3**).

Based on the data presented in the current study and a literature analysis, we can formulate a generalisable free energy model that depicts the ribosome-induced changes in free energies for unfolded, intermediate and native states. While the NC is constrained within the exit tunnel, folding is disfavoured due to the limited space available for folding and this is irrespective of NC net charge^13,33,41^. Upon complete emergence of (sub)domains from the tunnel, negatively and positively charged NCs diverge in their *de novo* folding behaviour (**Fig. 5d**). For negatively charged NCs, all states are destabilised by ribosome tethering, particularly the native state due to the unfavourable electrostatic potential energy contribution. The unfolded state is structurally expanded (due to electrostatic repulsion and lack of ribosome interactions throughout the sequence, **Fig. S6**) and entropically destabilised due to increased solvation^7^ relative to being off the ribosome. Intermediates are the least destabilised, thereby rationalising their higher populations on the ribosome compared to in isolation (**Fig. 5d**). Conversely, positively charged NCs do not populate stabilised, persistent coTF intermediates (or at least not more stable than in isolation)^9^. Their native states are either unaffected or stabilised by favourable electrostatic interactions with the ribosome^9,18,20^ compared to in isolation (**Fig. 5d**). Their unfolded conformations also experience long-range, attractive ribosome-NC interactions that result in a compaction and entropic stabilisation (**Figs. 5d, S6a-c** and ref.^9^). For all NCs, charge distribution across the surface of the folded state (**Fig. 3**) and throughout the sequence (**Figs. 4, S10, Table S2**) additionally modulates the magnitude of these effects for structured and unfolded states, respectively.

## Discussion

Multiple studies have observed the formation of natively folded structures to be less favourable on compared to off the ribosome^7,9,13,18–20,26,42^, but a general underlying model has been absent. We have previously shown that the ribosome can destabilise the unfolded state entropically and the native state enthalpically on the ribosome^7^, with both effects decreasing at longer linker length. In this work, we used NMR and simulations to develop a general conceptual framework that rationalise the NC length- and net charge-dependent coTF thermodynamics and pathways. Our data show that negatively charged NCs are destabilised when partially or fully folded due to an unfavourable (repulsive) contribution when tethered to the ribosome (**Figs. 3-4**): a negatively charged protein domain is less favoured near the negatively charged ribosome surface compared to in aqueous solution. These effects underpin the population of coTF intermediates not observed or less populated off the ribosome for several negatively charged proteins^7,11,13–17^ (**Fig. 5**). Indeed, ∼50% of all proteins carry a net negative charge suggesting this phenomenon is widespread^43^.

Our NMR and MD investigations of FLN5 as a model system also show that the ribosome does not induce structural and dynamic alterations in folded NC structure on the ribosome (**Fig. 1**). These findings align with previous studies that found NCs to become binding-competent for their native ligands during translation with similar binding affinities^10,17,20,44^. Alternative modes of destabilisation including ribosome interactions, entropic effects from protein dynamics or disordered linkers, or a reduced hydrophobic effect near the ribosome appear to contribute minimally (**Figs. 1-2, S4-S5**). The electrostatic model and its long-range nature also rationalise why coTF thermodynamics remain distinct from isolated polypeptides even at long linker lengths (>80 amino acids from the ribosome surface)^13^.

In line with our model (**Fig. 6**) and the net negative charge of the ribosome^45^, positively charged NCs (∼40% of the proteome^43^) are less susceptible to destabilising effects (**Table S2**). Instead, these NCs tend to have favourable interactions with the ribosome surface in both the unfolded and folded state, the balance of which determine their folding equilibria without populating stable coTF intermediates^9,18,20^. Additionally, long-range ribosome-NC interactions reduce the radius and solvent-accessible surface area of positively charged unfolded states (**Fig. S6a-c**), as observed in the compaction of unfolded T4 lysozyme on the ribosome^9^. The effects of charge on the unfolded state are thus entropically destabilising for acidic but stabilising for basic protein sequences due to their solvation and desolvation, respectively (**Figs. 5d, 6**). Positive charge consequently appears to suppress coTF intermediates being stabilised relative to other states during translation^9^. The ribosome can therefore act either as a holdase or foldase depending on the net charge and charge distribution of the NC.

**Figure 6.**
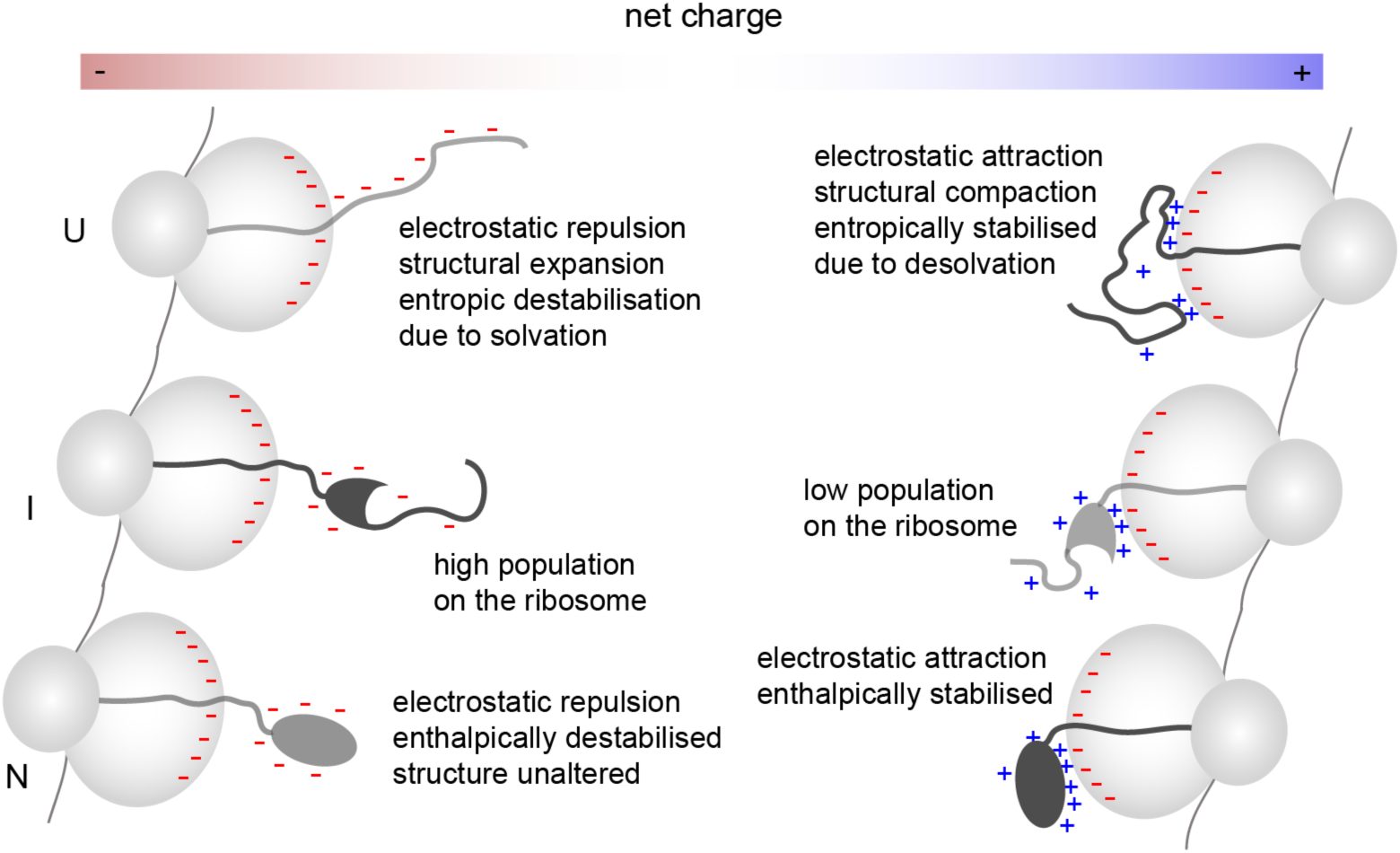
Net charge determines folding thermodynamics on the ribosome. The opacity of NC conformations is proportional to their generally observed population on the ribosome. Charge distribution throughout the sequence and folded surface further modulate the magnitude of these effects for unfolded and folded conformations, respectively.

Our findings imply that proteins with net negative charge are more prone to, at least partially in the case of coTF intermediates, exposing hydrophobic segments due to the delay in folding to the native state for up to ∼80 amino acids of biosynthesis^13^. Conformations with increased hydrophobic surface area are known to be crucial species in aberrant protein interactions and misfolding processes^46^. Their prolonged exposure during biosynthesis may make them more likely to misfold co-translationally, in contrast to positively charged sequences that can be sequestered by the ribosome^9,18,40,47^ and molecular chaperones^48,49^. Indeed, proteins that have been found to misfold co-translationally such as AAT^27^, CFTR NBD1^29,50^, calerythrin^12^, and EF-G^51^ all have a net negative charge, whereas T4 lysozyme (positively charged) is misfolding-prone off but not on the ribosome^9^. Relatedly, mature ribosomal proteins are highly stable as part of the ribosome complex but are prone to aggregation and degradation in isolation^52–54^.

Altogether, our study shows that long-range electrostatic forces shape coTF landscapes to modulate folding pathways and outcomes^7,8^. This model of protein folding thermodynamics on the ribosome may also well extend to proteins in other cellular environments, including coordination with co- or post-translational chaperones^16,17,55–57^ and proteins near anionic membranes, where similar long-range electrostatic effects may modulate protein free energy landscapes.

## Methods

### Sample production

We used standard, site-directed mutagenesis to introduce all mutations. Production of isotopically labelled (^1^H, ^15^N, ^19^F) protein samples of FLN5 and HRAS have been previously described^7,13,18^. ^19^F-labelled FLN5 RNC samples were also produced following previous protocols^13,18^.

### NMR spectroscopy

Residual dipolar couplings (RDCs) were measured with a 500 µM uniformly ^13^C,^15^N-labeled FLN5 sample in Tico buffer (10 mM HEPES, 30mM NH_4_Cl, 12mM MgCl_2_, 2mM β-merceptoethanol, pH 7.5) supplemented with 10% (v/v) D_2_O and 0.001% sodium trimethylsilylpropanesulfonate (DSS), at 298K on an 800 MHz Bruker Avance III HD spectrometer equipped with a TCI cryoprobe. After recording isotropic measurements, the sample was then partially aligned by the addition of liquid crystalline medium composed of C_12_E_5_ PEG and hexanol^58^. Hexanol was added stepwise until the phase transition was achieved, and alignment was verified by monitoring the splitting of the ^2^H resonance from the residual water signal (23.5 Hz). ^1^H-^15^N residual dipolar couplings were measured with a ^1^H,^15^N HSQC using the InPhase-AntiPhase (IPAP) approach^59^ (1 s recovery delay, 8 scans per increment and 197 ms acquisition time in the ^15^N dimension).

Backbone amide relaxation parameters^60–62^ were measured on a 280 µM uniformly ^15^N-labelled FLN5 sample in Tico buffer at 298K on four Bruker Avance III (500 MHz) or III HD (600, 700, 950 MHz) spectrometers all equipped with TCl/QCI-P cryoprobes. ^15^N-*R*_1_ values were measured from 2D spectra, recorded with 8 relaxation delays (500 and 700 MHz datasets: 40, 80, 240, 400, 560, 800, 1200, 1600 ms, 600 and 950 MHz datasets: 10, 50, 100, 200, 400, 600, 900 1200 ms). ^15^N-*R*_2_ values were measured from 2D spectra recorded, for the 600 and 950 MHz datasets with 8 relaxation delays (8, 16, 24, 32, 48, 64, 96, 152 ms). For the 500 and 600 MHz datasets spectra were recorded with an *R*_1,rho_ experiment with 8 relaxation delays (1, 5, 10, 20, 40, 70, 110, 150 ms) and ^15^N-*R*_2_ rates were subsequently calculated. Spectra were processed and analysed using nmrPipe^63^. Peak intensities were fitted to two-parameter decay curves *I*(*t*) = *I*0e^-*Rt*^, where *I* is the magnetisation intensity, *t* is the relaxation delay and *R* is the relaxation rate. ^1^H-^15^N NOE measurements were collected by recording 2D spectra with and without saturation. ModelFree^64^ was used to obtain internal dynamics and global tumbling parameters with the model-free approach^65^.

1D ^19^F NMR experiments were acquired at 298 K on a 500 MHz Bruker Avance III spectrometer, equipped with TCl cryoprobes, with TopSpin3.5pl2. An acquisition time and recycle delay of 350 ms and 1.5 s, respectively, were used. As previously described, we recorded experiments successively and only summed spectra once the stability of RNC samples has been ensured by examining the spectra during course of the experiment^7,13^. Data were analysed and processed with nmrPipe^63^, CCPN Analysis^66^ and MATLAB (R2014b, The MathWorks Inc.) as previously described^13^. Lorentzian lineshapes were fit (see Code Availability) to determine the linewidths and integrals (populations), and errors were obtained as the standard error of the mean from bootstrapping iterations^7,13^. Samples were measured in Tico buffer supplemented with 10% D_2_O (v/v) and 0.001% DSS. Folding free energies were calculated directly from the fractional populations of two states, for example A and B, using ΔG = -RTln(A/B). Linewidths were converted to rotational correlation times of FLN5 using a previously established empirical relationship for FLN5^7^.

### MD simulations and computational analyses

#### Calculation of FLN5 surface electrostatic potentials

We used PyMOL 2 (Schrödiner, LLC) to introduce mutations in *in silico* into FLN5 starting from the crystal structure (PDB 1QFH, chain A, residues 646-750)^67^. Surface electrostatic potentials were then calculated using the PDB2PQR^68^ and APBS^69^ plugins in PyMOL.

#### All-atom MD simulations of isolated FLN5

We used GROMACS^70^ v2023 to prepare and run all MD simulations described. Simulations were initiated from the crystal structure of FLN5 (PDB 1QFH, chain A, residues 646-750)^67^, and we additionally built the N-terminal MHHHHHHAS tag as an extended chain in PyMOL. The protein was then centred in a dodecahedral box with a minimum distance of 2 nm from the box edge (box volume ∼916 nm^3^), solvated and neutralised. A final MgCl_2_ concentration of 12 mM was added and the resulting box of FLN5 contained 29,614 water molecules. We used the DES-Amber protein force field^71^ and TIP4P-D water model^72^. The system was energy minimised using steepest decent and equilibrated with the leap-frog integrator and a 2 fs timestep with bonds involving hydrogen constrained using the LINCS algorithm ^73^. Short-range distance cut-offs of 0.9 and 1.0 nm were used for van der Waals and electrostatic interactions, respectively. Long-range electrostatics were computed using the Particle Mesh Ewald (PME)^74^ with cubic interpolation and a grid spacing of 0.125 nm. Temperature coupling was done using the velocity rescaling algorithm^75^ with a time constant of 0.1 ps. An initial equilibration in the NVT ensemble was performed at 298 K for 500 ps using position restraints on all heavy atoms with a force constant of 1000 kJ mol^-1^ nm^2^. The box volume was then equilibrated with a NPT simulation for 500 ps using the Berendsen barostat^76^ at a pressure of 1 bar, compressibility of 4.5×10^-5^ bar^-1^ and coupling constant of 2 ps. Position restraints were then removed for production simulations and pressure coupling was switched to the Parrinello-Rahman algorithm^77^. We employed hydrogen mass repartitioning^78^ and used a time step of 4 fs for production simulations. Eight independent simulations were run for 2 μs each. Protein coordinates were saved every 2 ps and the whole system coordinates every 100 ps.

#### All-atom MD simulations of FLN5 RNCs

We performed all-atom MD simulations of folded FLN5 linked to the ribosome by two different linker lengths, FLN5+34 and FLN5+47. We used the same model of the ribosome (containing atoms near the exit tunnel and surrounding surface at the exit vestibule) and structures of the FLN6-SecM linker as in prior work^7,18^ (Mitropoulou *et al.*, in preparation). His-tagged, folded FLN5 was covalently linked to the linker models to generate initial starting structures of the complex. We placed the complexes in a rhombic dodecahedron box with a diameter of 19.621 nm and volume of 14,001 nm^3^. The complexes were solvated and neutralised with Mg^2+^ ions, resulting in 1,927,565 and 1,927,515 atoms for FLN5+34 and FLN5+47 with a four-point water model, respectively. We used the DES-Amber protein^71^ and nucleic acid^79^ combined with the TIP4P-D water model^72^. Simulations were also run with a scaled version of the force field where all protein-protein and protein-nucleic acid Lennard-Jones interactions were scaled by a factor of 0.9 (LJ09). Coupling and constraint algorithms used were identical to the isolated protein simulations. After energy minimisation, we first progressively heated the system from 0 to 298 K during a 500 ps NVT simulation, followed by a 1 ns NVT equilibration at 298 K. The volume and density were equilibrated in a 1 ns NPT simulations at a pressure of 1 bar. All heavy atoms were position restrained (1000 kJ mol^-1^ nm^2^) during equilibration, and position restraints on the nascent chain molecule were then removed for production (except for the C-terminal residue at the peptidyl transferase centre to prevent release). Production simulations were run with hydrogen mass repartitioning and time step of 4 fs and initiated from different initial conformations/orientations on the ribosome obtained after an initial 100 ns of unbiased MD. Eight simulations of 2 μs were then performed from different starting structures and protein/RNA and system coordinates were saved every 2 ps and 200 ps, respectively.

#### Comparison of MD simulations with experimental data

RMSD and RMSF calculations were performed with respect to the crystal structure of FLN5 (PDB 1QFH) using the GROMACS tools gmx rms and gmx rmsf, respectively. Pairwise RMSD matrices were calculated using MDAnalysis^80^. All NMR experimental parameters were ensemble-averaged over all simulation trajectories. Chemical shifts were calculated using SHIFTX2^81^. Amide N-H bond vector residual dipolar couplings (RDCs) were calculated using the PALES software with the global molecular alignment method based on molecular shape and the infinite wall model^82^. RDCs were then globally scaled to optimise the agreement with experimental data. Agreement between calculated and experimental RDCs was calculated using the quality factor (Q-factor) metric^83^:

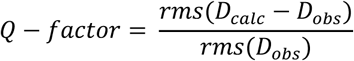

Amide N-H order parameters were calculated from the values of the internal correlation functions of respective N-H bond vectors at lag times corresponding to the experimental rotational correlation time of 7.7 ns^31^. Internal correlations functions were calculated after alignment of all MD snapshots to a reference structure using backbone heavy atoms of residues 646-750 to remove rotational and translational contributions to the correlation function. The ensemble averaged correlation functions were then calculated using a second-order Legendre polynomial^84^ where *θ* is the angle between unit vectors of an amide bond temporally separated by a lag time, t:

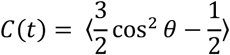

To compare the strength of ribosome-nascent chain (NC) interactions observed in MD simulations with experiments, we quantified the rotational correlation time (τ_C_) of FLN5 in the RNC simulations and normalised it against the value of the isolated protein (i.e., the ratio by which the ribosome reduces rotational tumbling) (Mitropoulou *et al.*, in preparation). These ratios have previously been measured experimentally for natively folded FLN5 by ^13^C NMR^31^ and ^19^F NMR^7,13^. We first aligned all MD snapshots against a reference structure using all backbone atoms in the β-strand region. The resulting rotation matrices were then applied to a set of 1,000 uniformly distributed orientation vectors in the unit sphere. The rotational correlation function was then calculated using a second-order Legendre polynomial where *α* is the angle between two FLN5 orientations separated by a lag time, τ:

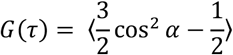

Using Scipy (optimize.curve_fit)^85^, these combined data from all 1,000 unit sphere vectors were used to determine τ_C_ by globally fitting *G*(τ) up to lag times of 50 ns to a single exponential decay for the isolated protein:

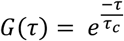

The mean and standard error of the mean were determined from eight independent simulations lasting 2 μs. For RNCs, we fitted the data to a double exponential function to the average (weighted by the standard deviation) from all 1,000 unit spheres where p_1_ and p_2_ are the amplitudes (populations) of the two decays and S^2^_tether_ accounts for the restricted tumbling of FLN5 due to position restraints on the ribosome particle:

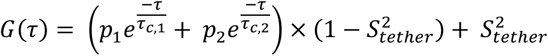

S^2^_tether_ has a value of 1 when FLN5 is rigidly bound to the ribosome surface and 0 if FLN5 tumbles independently of the ribosome. Thus, we set S^2^_tether_ to 0 when the correlation functions reach 0. Double exponential fits were done up to lag times of 1,000 ns. For each independent MD trajectory of FLN5 RNCs, the ensemble averaged τ_C_ of FLN5 was then calculated as:

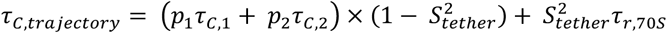

τ_r,70S_ is the value of the ribosome correlation time. To estimate this, we used the experimental value (∼ 3,000 ns at 298 K in water)^86^ and scaled it by 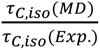, where τ_C,iso_ is the rotational correlation time of FLN5 in isolation. This scaling factor accounts for the factor change of diffusion and tumbling constants due to the water model against the known tumbling time of FLN5. The final average and standard error of the mean of the FLN5 RNCs was then calculated from eight independent simulations and values of τ_C,trajectory_.

To orthogonally assess and compare ribosome-NC interactions between different RNC simulations we calculated the residue-specific interaction probabilities using a contact definition of < 10 Å considering Cα protein atoms and P, N3 and C4’ RNA atoms. We also calculated the minimum distance between the ribosome surface and FLN5 centre of mass and minimum distance between the ribosome surface and any FLN5 atom using the GROMACS tools gmx pairdist and gmx mindist, respectively. With gmx mindist, we also calculated the fraction of FLN5 atoms that are within 1.0 nm of the ribosome.

#### Calculation of thermodynamic parameters of folded FLN5

We predicted the thermodynamic differences between isolated and ribosome-bound protein from the MD simulations. To obtain the effects of solvation, we first used GROMACS gmx rdf to calculate the radial distribution function (distance between the centres of mass of water molecules and the protein surface) to identify the hydration layer (first two hydration shells exhibiting sharp peaks in the radial distribution function). Based on this analysis, we selected a distance cut-off of 0.35 nm to calculate the number of solvation layer water molecules of FLN5. Entropic and enthalpic changes due to solvation were estimated separately based on the solvent-accessible surface area (SASA) of residues 646-749. We defined polar and apolar surface areas by atoms having an absolute partial charge of more than or less than 0.3, respectively. Empirical parameters relating SASA changes to thermodynamics were used^87,88^, as previously described^7^. Conformational entropy changes were calculated from backbone and sidechain probability distributions as previously described^7,89^. The total bonded potential energies (bond, angles, dihedrals, improper dihedrals) were obtained from the DES-Amber force field parameters using gmx mdrun -rerun, including all bonded terms of residues 646-749. The nonbonded potential energy of FLN5 was calculated by defining FLN5 (residues 646-749) as a group and summing all intramolecular (1-4 and short-range terms) and intermolecular (short-range terms between FLN5 and the rest of the simulation system) nonbonded terms. A distance cut-off of 3.5 nm was used to compute the short-range term to avoid neglecting contributions from long-range interactions. All potential energy rerun calculations were performed with GROMACS 2020^70^. The thermodynamic quantities were averaged over all eight trajectories of 2 μs and the standard error of the mean was calculated from the eight individual trajectories. For the conformational entropy analysis, we performed a blocking analysis assessing the entropy differences and errors as a function of block size since conformational entropy depends on the total amount of sampling up to block sizes of ∼8 μs. Thus, we calculated the error from two halves of 8 μs.

#### Water entropy calculations

The molar entropies of water as a function of the distance from the ribosome surface were calculated with the two-phase thermodynamic (2PT) method^90^ and DoSPT code (https://dospt.org/index.php/DoSPT)^91^. We used the CHARMM36m force field with the CHARMM-modified TIP3P water model with increased water dispersion interactions (water hydrogen ε parameter of −0.1 kcal mol^-1^)^92^. The TIP3P family of water models has been shown to most accurately reproduce the standard molar entropy of water compared to experiments^90^. We used a partial ribosome model employed in previous simulation work of an unfolded FLN5 RNC^7^, and ran five independent simulations of 20 ps in the NVT ensemble using position restraints on ribosome heavy atoms (1000 kJ mol^-1^ nm^2^). Velocity and coordinate data were saved every 4 fs and a time step of 2 fs with LINCS constraints on all hydrogen bonds was used. The velocity verlet (md-vv) integration algorithm was used to obtain accurate velocity data. The remaining simulation parameters for this force field were previously described for an unfolded FLN5 RNC^7^. The system configuration was initially equilibrated for 20 ns in the NPT ensemble prior to production data collection for 2PT calculations. Water molecules remaining within a defined distance region from the ribosome surface for the entire 20 ps simulation were analysed and the effective volume was estimated based on the average water molecule volume obtained previously under identical conditions^7^. Local water entropies are reported as the average and standard error of the mean from five independent simulations.

#### Coarse-grained MD simulations and linker entropy calculations

Coarse-grained MD simulations of disordered C-terminal linkers and FLN5/HRAS proteins extended by their C-terminal linkers (built in an extended conformation in PyMOL) were performed using the Martini3 force field^93^ and GROMACS v2021^70^. The Lennard-Jones interaction parameters between all protein beads were scaled by a factor of 0.88, which has been shown to lead to improved agreement for disordered and multidomain proteins with SAXS and NMR data^94^. All simulations were performed at an ionic strength of 50 mM using NaCl. The disordered linkers were modelled in an all-coil conformation without employing any sidechain (scfix) or contact (elastic network) restraints on their structure and dihedrals. The FLN5 and HRAS systems with their linkers were modelled using secondary structure assignments obtained with the DSSP algorithm^95^. An elastic network model was applied with Martinize2^96^. All backbone bead pairs within 0.9 nm were restrained by a harmonic potential of 700 kJ mol^-1^ nm^-2^. Angle and dihedral restraints between backbone and sidechain beads were applied with the -scfix flag based on the input structures. We used the FLN5 crystal structure (PDB ID 1QFH^67^). A solvated input conformation was energy minimised with the steepest descent algorithm. The systems (FLN5 with two linkers of 5 and 15 amino acids) were then equilibrated in the NPT ensemble for 10 ns using a 2 fs timestep. The temperature was set to 298 K and maintained using velocity rescaling^75^ and the pressure of 1 bar was kept constant using the Parrinello-Rahman method^77^. The timestep was then increased to 20 fs and an additional 1 μs of MD was performed to equilibrate the systems. A final production simulation of 50 μs was then run for all linker peptides and proteins, saving conformations every 0.1 ns for analysis.

To predict changes in stability based on these coarse-grained simulations, we considered the conformational entropy of the C-terminal linker and changes in solvent-accessible surface area (SASA). SASA values were calculated using the gmx sasa tool, recommended van der Waals radii for Martini3 beads^93^ and a probe radius of 0.191 nm. For conformational entropy calculations we back-mapped all coarse-grained snapshots to all-atom resolution using cg2all^97^ (https://github.com/huhlim/cg2all). Entropies were then calculated as previously described in ref.^7^. Errors were estimated from a blocking analysis.

## Data/code availability

The NMR chemical shift assignment of FLN5 has been previously deposited in the BMRB under the entry code 51075. The structural ensembles and MD trajectories of folded FLN5 on and off the ribosome will be deposited on Zenodo. Code used to fit the ^19^F NMR spectra are available on Github (https://github.com/shschan/NMR-fit).

## Acknowledgements

This work was supported by a Wellcome Trust Investigator Award (to J.C., 206409/Z/17/Z). We acknowledge use of the UCL Biomolecular NMR Centre and the MRC Biomedical NMR Centre at the Francis Crick Institute. The Francis Crick Institute receives its core funding from Cancer Research UK (FC001029), the UK Medical Research Council (FC001029) and the Wellcome Trust (FC001029). We acknowledge the use of computational resources provided by the Baskerville Tier 2 HPC service (https://www.baskerville.ac.uk/). Baskerville was funded by the EPSRC and UKRI through the World Class Labs scheme (EP/T022221/1) and the Digital Research Infrastructure programme (EP/W032244/1) and is operated by Advanced Research Computing at the University of Birmingham. This project also made use of time on HPC resources on Archer2 (ARCHER2 UK National Supercomputing service, https://www.archer2.ac.uk) granted via the UK High-End Computing Consortium for Biomolecular Simulation, HECBioSim (http://hecbiosim.ac.uk), supported by EPSRC (grant no. EP/R029407/1 and EP/X035603/1). We additionally acknowledge the use of the UCL Myriad High Performance Computing Facility (Myriad@UCL) and associated support services in the completion of this work.

