## Supplementary Information for "Long-range electrostatic forces govern how proteins fold on the ribosome"

### Supplementary text

#### *MD simulations of folded FLN5 on and off the ribosome*

First, two state-of-the-art force fields, CHARMM36m[92] and DES-Amber[71,72,79], were compared to ensure accurate structural insights are obtained. These were evaluated against NMR backbone chemical shifts[31], residual dipolar couplings and relaxation measurements converted to amide order parameters for isolated FLN5 protein (see Methods), as well as measurements of ribosome interactions of FLN5 RNCs detected via an increase in the domain's rotational correlation time (see Methods). With the CHARMM36m[92] (C36m) force field, the simulations of the isolated protein (Mitropoulou *et al.*, in preparation) exhibit an average RMSD of 2.1 ± 0.1 Å from the crystal structure (**Fig. S1a-b**), whereas with DES-Amber-3.20[71,79] the simulations agree more closely with the experimental structure with an RMSD of 1.5 ± 0.0 Å (**Fig. S1a-b**). When compared to solution NMR data, we find that DES-Amber also agrees more closely with residual dipolar couplings (**Fig. S1c**), backbone chemical shifts (**Fig. S1d-f**) and amide order parameters (**Fig. S1g**) compared to C36m. The latter reveals that C36m predicts the C-terminus and loop regions to be too dynamic compared to NMR data, and DES-Amber results in reduced backbone dynamics that agree better with experiments (**Fig. S1h**).

We encouragingly find that DES-Amber also results in a stable FLN5 structure on the ribosome with an RMSD of 1.5 ± 0.0 and 1.6 ± 0.0 Å at linker lengths of FLN5+34 and FLN5+47, respectively (**Fig. S2a-b**). At a linker length of 47 amino acids (FLN5+47), DES-Amber simulations show good agreement with the experimentally measured rotational correlation time, similar to our previous results with C36m with an increased protein/RNA-water dispersion interaction (C36m+W) (Mitropoulou *et al.*, in preparation), suggesting that ribosome-nascent chain interactions are accurately captured (**Figs. S2c-S3**). FLN5+47 simulations with DES-Amber are thus analysed in depth. In line with NMR-observed resonances, ribosome-nascent chain interactions are highly transient (0-20% of ribosome contacts depending on the region of FLN5, **Fig. S2e**), and contacts involve most structural elements surrounding the exit tunnel (**Fig. S2f**). Interactions are observed in loop regions of residues 665-750 and 720-725 as well as the C-terminus (residues 745-750), which together make up the C-terminal face of FLN5 closest to the tether (linker) to the ribosome (**Fig. S2e**).

Conversely, at the shorter linker length of FLN5+34, both C36m and DES-Amber significantly overestimate the rotational correlation time of FLN5 due to stronger than experimentally quantified interactions with the ribosome (**Figs. S2c-S3**). The linker in the ribosome exit tunnel remains in an extended conformation in all simulations and the linker dimensions are in good agreement with distances obtained from cryo-EM analyses (Mitropoulou *et al.*, in preparation) (**Fig. S2d**).

We further attempted to improve the agreement with for shorter RNCs by reducing protein-protein and protein-RNA Lennard-Jones interactions of the DES-Amber force field by 10% ('LJ09', see Methods).

Although the protein remains completely folded in all simulations, this results in a slightly worse agreement with the crystal structure and NMR data compared to the original DES-Amber force field (**Fig. S1**). At FLN5+34 the agreement with the rotational correlation time has improved slightly but is still considerably overestimated (**Figs. S2c-S3**). In the main text, our analyses thus describe results obtained with the original DES-Amber-3.20 force field comparing the isolated protein with the FLN5+47 RNC, where on balance we achieve the best agreement with NMR data of the isolated protein and ribosome interactions. Analyses of the LJ09 are shown in the Supplementary Information (**Figs. S1-S4**).

#### *Analysis of water dynamics and entropy near the charged ribosome surface*

Using our simulation models, we further assessed whether alternative mechanisms of destabilisation may explain coTF thermodynamics. A previous theoretical study suggested that proteins may be destabilised near the ribosome due to a reduced magnitude of the hydrophobic effect because water molecules near the charged ribosome surface have lower entropy[22]. Consequently, upon protein folding and release of water from the hydration layer into bulk solution, the entropic gain would be lower. Water ordering and entropy are expected to be affected close to the ribosome surface as, for example, within the tunnel and vestibule[22]. Indeed, we also find that water entropies are only attenuated within ~1 nm of the ribosome surface using MD and two-phase thermodynamics calculations (see Methods, **Fig. S5a-b**), in agreement with theoretical calculations showing reduced water entropies near lipid bilayers up to ~1 nm from the surface[98].

Thus, these effects may play a role for smaller proteins, such as helical bundles that can form even within the tunnel and vestibule[15,99-101]. Larger proteins like FLN5, however, are expected to form their hydrophobic core at distances beyond 1 nm of the ribosome surface. Indeed, at FLN5+47 the centre of mass of FLN5 is ~2-2.5 nm away from the ribosome on average, and only 3-8% of FLN5 atoms are within 1 nm of the ribosome surface (**Fig. S5c-d**). This is even more the case for unfolded nascent chain conformations, where distances of multiple tens of nanometres from the ribosome surface are sampled[7,40]. Modulated water dynamics near the charged ribosome surface thus cannot account for the destabilisation, in line with this destabilisation being a long-range effect lasting over at least 80 amino acids of FLN5 biosynthesis[13]. Recent measurements of the folding entropy on the ribosome, which revealed that the entropic penalty was lower on compared to off the ribosome, also strongly indicate that the hydrophobic is, in fact, stronger and not weaker on the ribosome[7].

### Supplementary figures

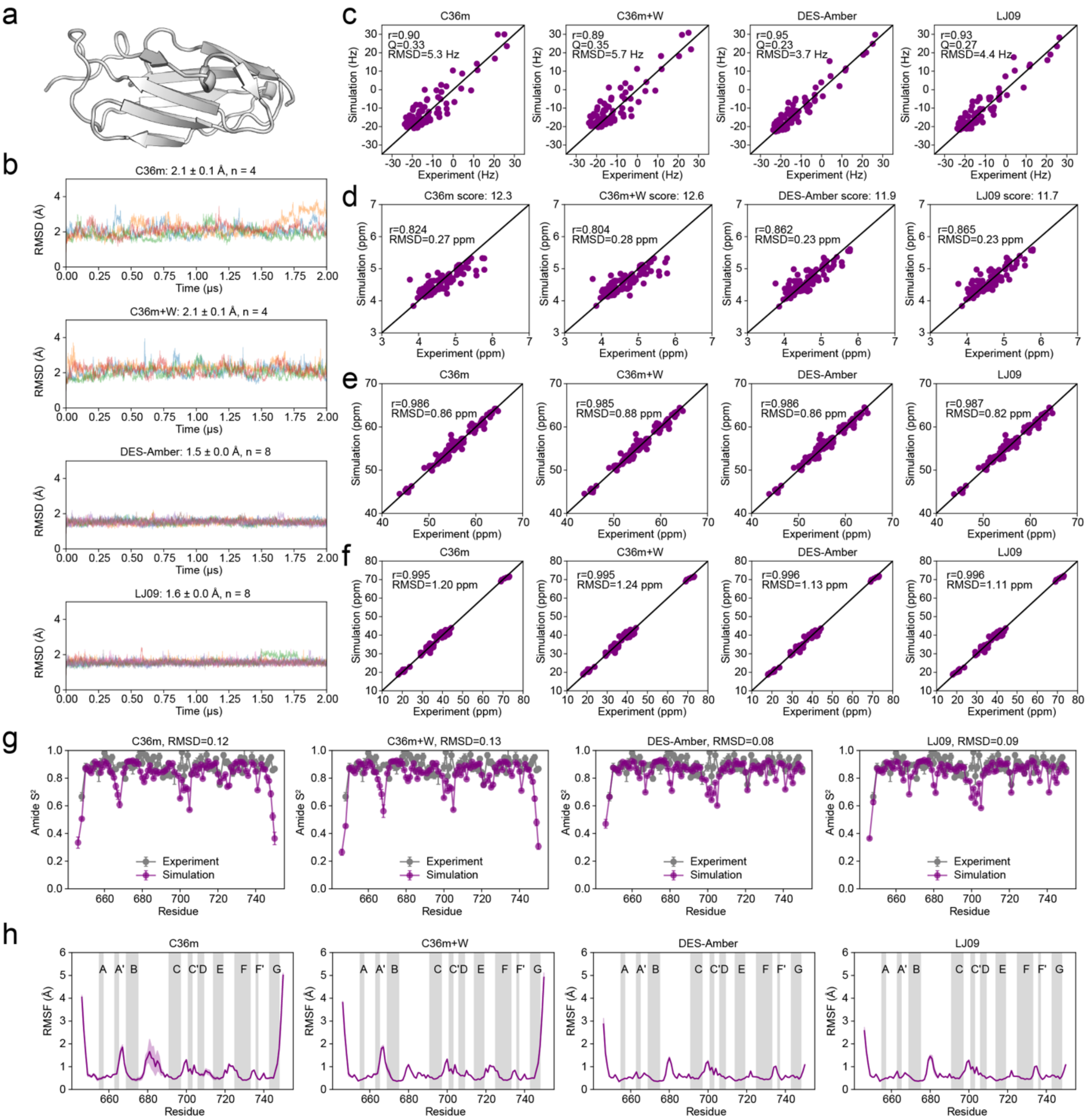


**Figure S1.** Comparison of isolated FLN5 MD simulations with experimental data. **a.** Crystal structure (PDB 1QFH[67]) of FLN5 residues 646-750. **b.** All-atom root mean square deviation (RMSD) of residues 646-750 relative to the crystal structure observed in MD simulations with different force fields (mean ± s.e.m.). **c.** Experimental and calculated residual dipolar couplings (RDCs) of amide N-H bonds with different force fields. **d-f.** Experimental and calculated backbone (Hα, d; Cα, e; Cβ, f) NMR chemical shifts with different force fields. The score in panel d is the sum of nucleus specific RMSDs normalised by their average prediction error[81] from five nuclei (N, HN, Cα, Hα, and Cβ). **g.** Amide order parameters determined from NMR relaxation experiments and calculated from MD simulations with different force fields (mean ± s.e.m.). **h.** Backbone root mean squared fluctuations (RMSF) calculated at the Cα atom with different force fields (mean ± s.e.m.).

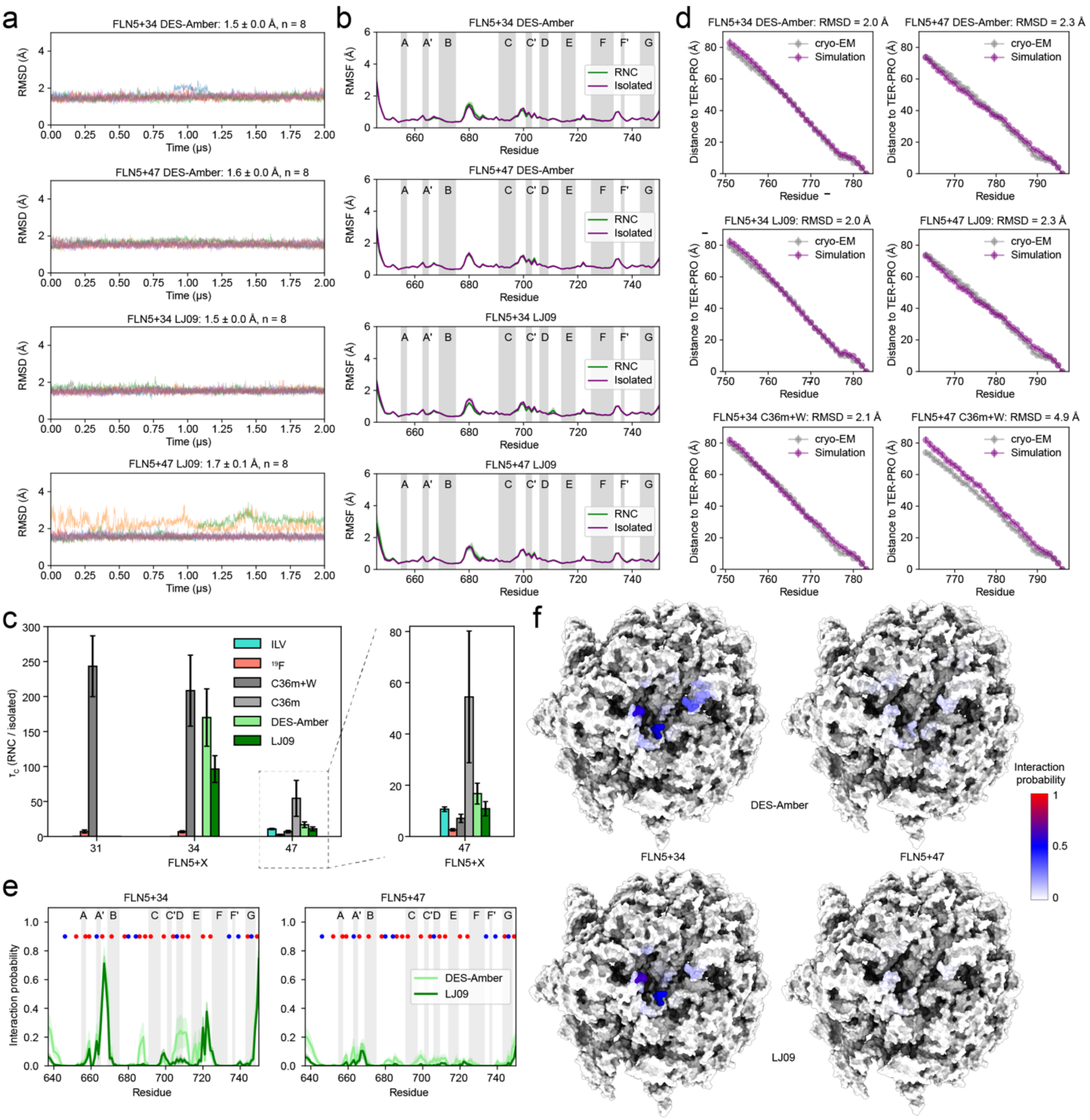


**Figure S2.** Comparison of FLN5 RNC MD simulations with experimental data and characterisation of ribosome interactions. **a.** All-atom root mean square deviation (RMSD) of residues 646-750 relative to the crystal structure observed in MD simulations of FLN5+34 and FLN5+47 with DES-Amber 3.20 and LJ09 (mean ± s.e.m.). **b.** Backbone root mean squared fluctuations (RMSF) calculated at the $C\alpha$ atom with DES-Amber 3.20 and LJ09 from MD simulations of FLN5+34 and FLN5+47 compared with the isolated protein (mean ± s.e.m.). **c.** Comparison of experimentally measured and simulated (MD) FLN5 rotational correlation times, $\tau_C$, normalised by their respective values of the isolated protein. Experimental values were previously obtained by $^{13}$C/ILV NMR[31] and $^{19}$F NMR[7,13]. MD values obtained with the C36m force fields were previously published (Mitropoulou *et al.*, in preparation). **d.** Effective linker lengths (comprised of the arrest-enhanced SecM stalling sequence and FLN6 linker) calculated as the average distance (mean ± s.d.) from the C-terminal proline at the PTC observed in cryo-EM structures (Mitropoulou *et al.*, in preparation) and MD simulations with different force fields. **e-f.**

Ribosome-nascent chain interaction probability calculated from MD simulations with DES-Amber 3.20 and LJ09 mapped along the sequence of FLN5 (mean ± s.e.m.) and on the ribosome surface.

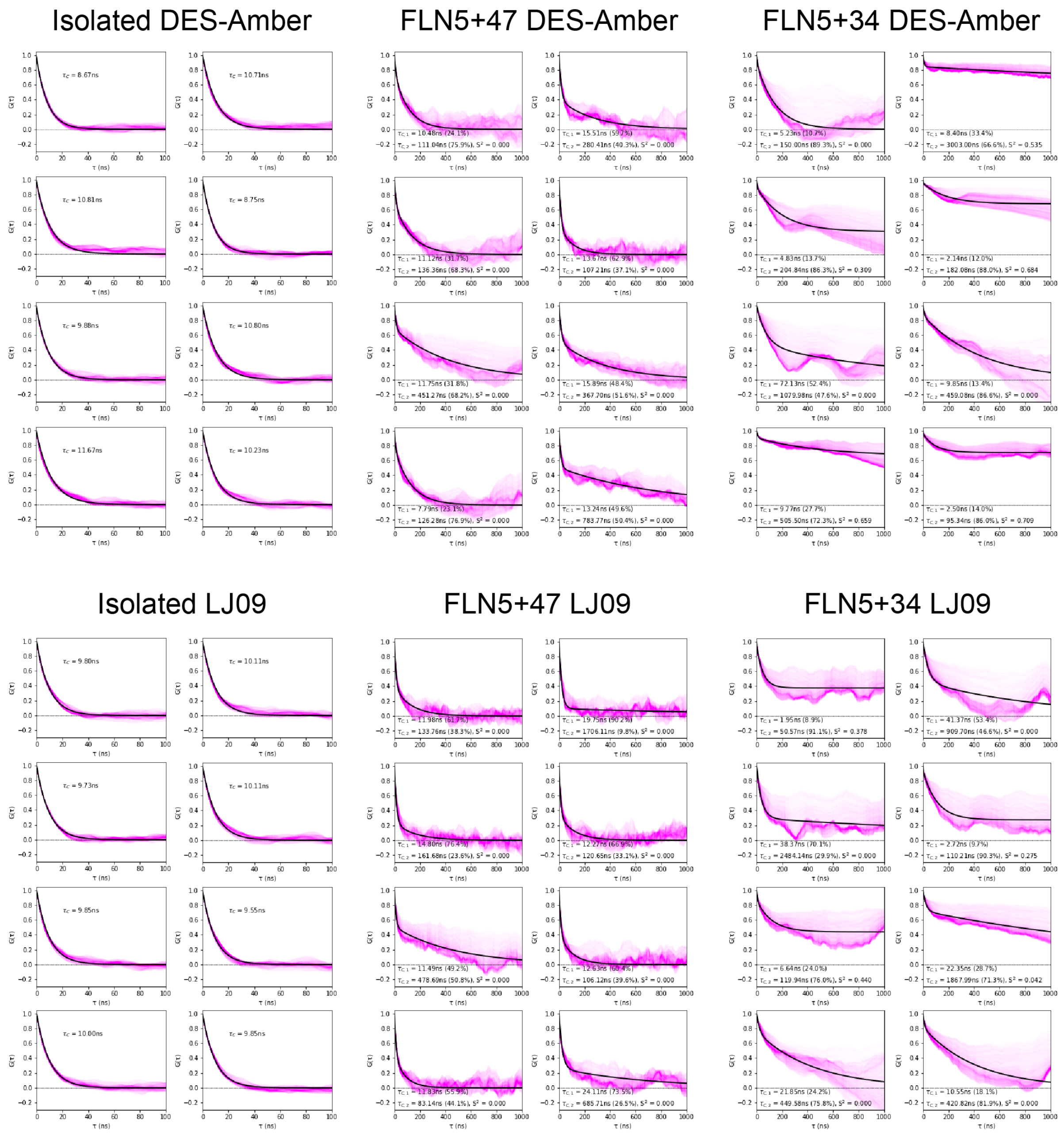


**Figure S3.** Fits of rotational correlation functions to all individual MD trajectories of FLN5 on and off the ribosome. The magenta lines correspond to all of the orientational vectors (1,000 randomly sampled in a unit sphere) and black lines correspond to the fits. The individually fitted rotational correlation times and amplitidues ($\tau_C$, %) and plateau values ($S^2$) are shown on each plot.

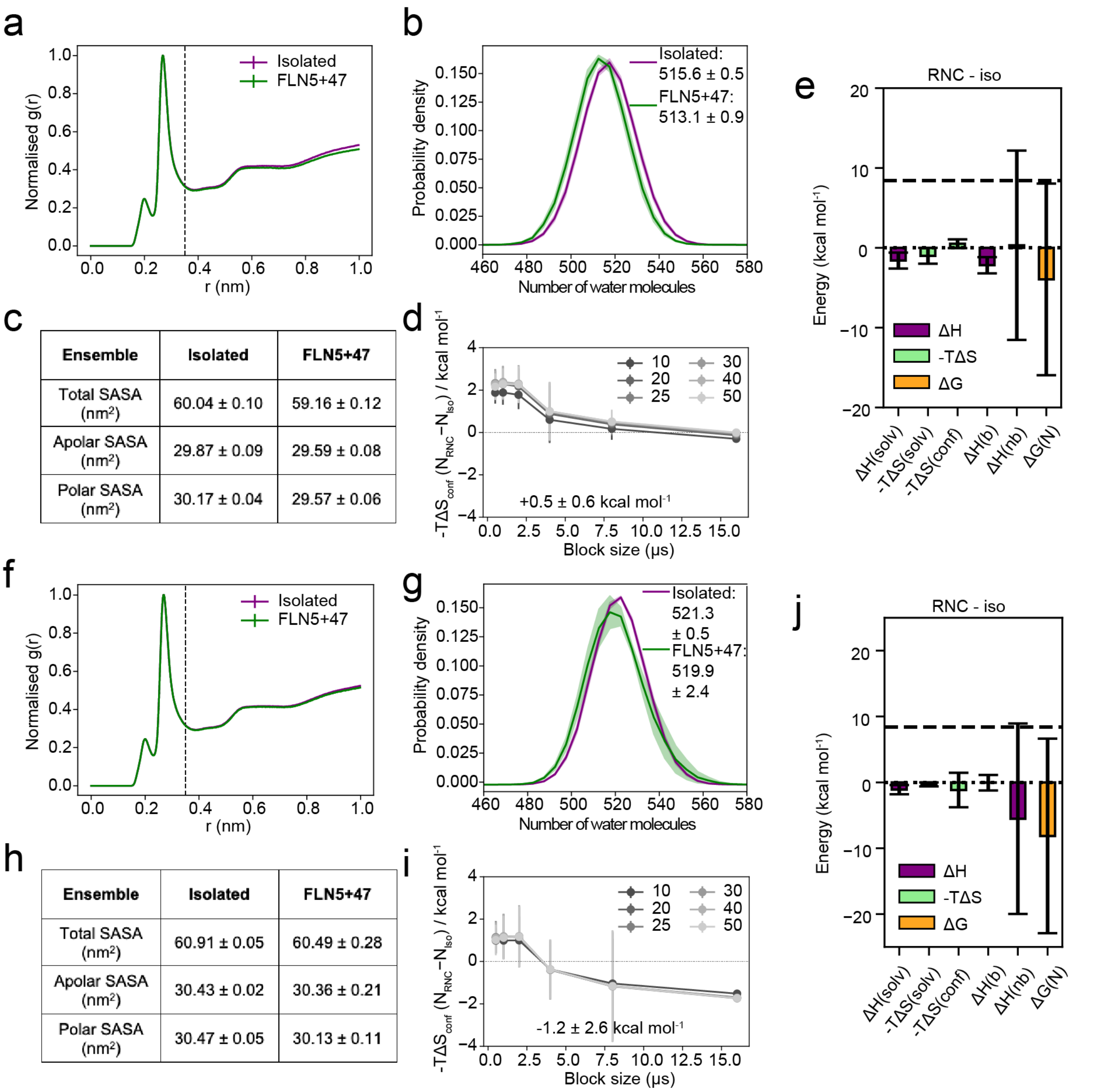


c

| Ensemble | Isolated | FLN5+47 |
|---|---|---|
| Total SASA (nm²) | 60.04 ± 0.10 | 59.16 ± 0.12 |
| Apolar SASA (nm²) | 29.87 ± 0.09 | 29.59 ± 0.08 |
| Polar SASA (nm²) | 30.17 ± 0.04 | 29.57 ± 0.06 |

h

| Ensemble | Isolated | FLN5+47 |
|---|---|---|
| Total SASA (nm²) | 60.91 ± 0.05 | 60.49 ± 0.28 |
| Apolar SASA (nm²) | 30.43 ± 0.02 | 30.36 ± 0.21 |
| Polar SASA (nm²) | 30.47 ± 0.05 | 30.13 ± 0.11 |

**Figure S4.** Solvation, dynamics, and energetics of folded FLN5+47 compared to isolated FLN5 calculated from MD simulations with DES-Amber 3.20 (**a-e**) and LJ09 (**f-j**). **a,f.** Radial distribution functions of the water centre of mass to protein distance (mean ± s.e.m.). The vertical line at 0.35 nm shows the cut-off used to define the hydration layer (first two hydration shells). **b,g.** Probability distributions and average of the number of water molecules in the hydration layer of FLN5 (mean ± s.e.m.). **c,h.** Average total, apolar and polar SASA calculated for FLN5 ensembles on and off the ribosome (mean ± s.e.m.). **d,i.** Difference in protein conformational entropy of FLN5 on and off the ribosome as a function of the number of bins used to discretise dihedral distributions (legend) and simulation/sampling block size of the entire, concatenated ensembles (mean ± s.e.m.). The final values (50 bins, 8 μs blocks) are shown on the plot. **e,j.** Bar plot summarising predicted energetic changes of folded FLN5 (RNC-isolated protein) from solvation, conformational entropy, bonded, and nonbonded potential energy (see Methods). The dashed, horizontal line represents the lower bound estimate of the native state destabilisation previously determined[7]. All values represent the mean ± s.e.m. from eight independent simulations of 2 μs. The s.e.m. of the conformational entropy term was calculated using a blocking analysis and a 8 μs block size (panels d,i).

a

| Water entropy at 298K (J mol$^{-1}$ K$^{-1}$) | |
|---|---|
| Experiment | 69.95 ± 0.03 |
| TIP3P literature | 68.49 ± 0.14 |
| C36m (mTIP3P) | 68.50 ± 0.04 |
| C36m+W (mTIP3P+W) | 68.50 ± 0.05 |

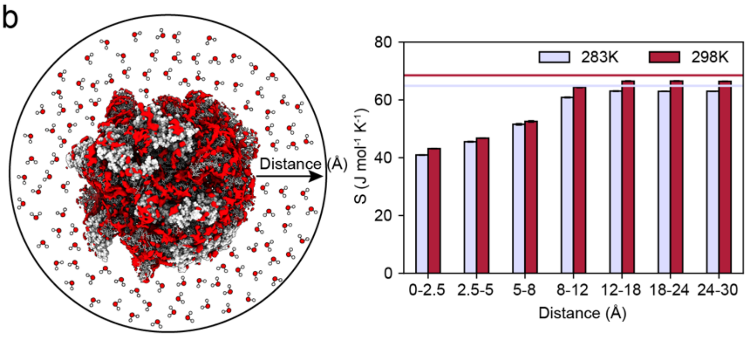


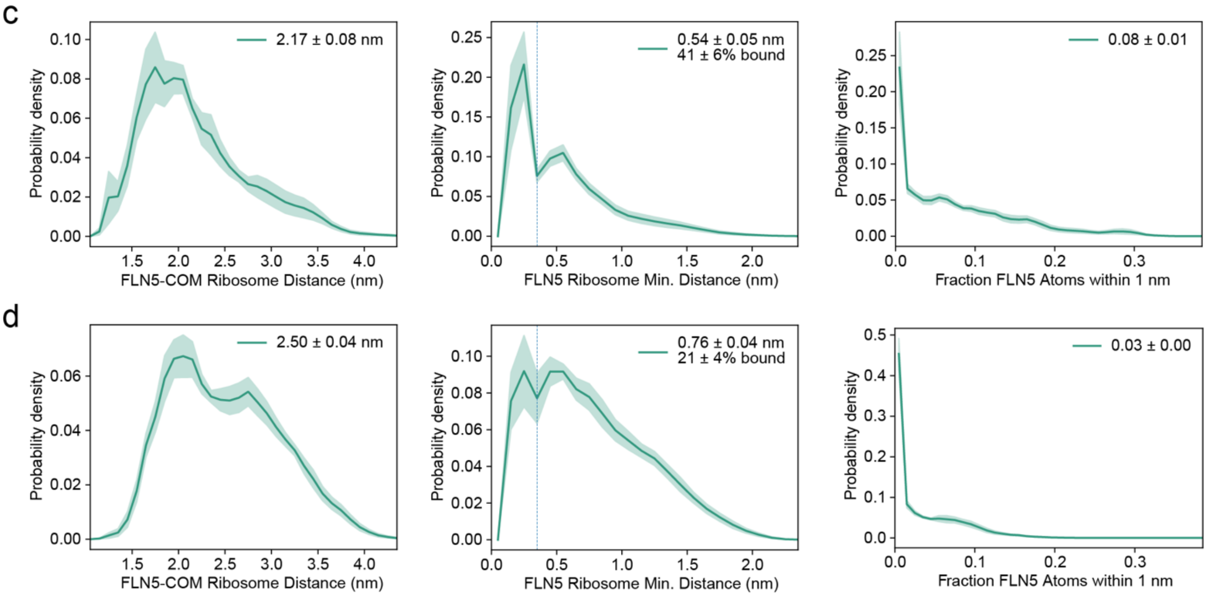


**Figure S5.** Water entropy near and distance of FLN5 from the ribosome surface. **a.** Molar entropies of water under standard conditions (298 K, 1 bar) obtained by experiments[102], the literature value for the TIP3P water model[90], and our previously obtained values with the CHARMM TIP3P water models (+W with an increased water dispersion strength; water hydrogen $\varepsilon$ = -0.1 kcal mol$^{-1}$) calculated for a pure water box[7]. **b.** Molar entropy of water as a function of distance from the ribosome surface calculated using the two-phase thermodynamics method at two temperatures. The horizontal lines correspond to the bulk values obtained from a pure water box at these temperatures. **c-d.** Probability distributions (mean ± s.e.m.) and ensemble averaged values of FLN5 (centre of mass, COM) ribosome minimum distances (left), FLN5 ribosome minimum distances (middle) and fraction of FLN5 atoms within 1 nm of the ribosome. These values were calculated for MD simulations with the DES-Amber 3.20 (panel c) and LJ09 (panel d) force fields.

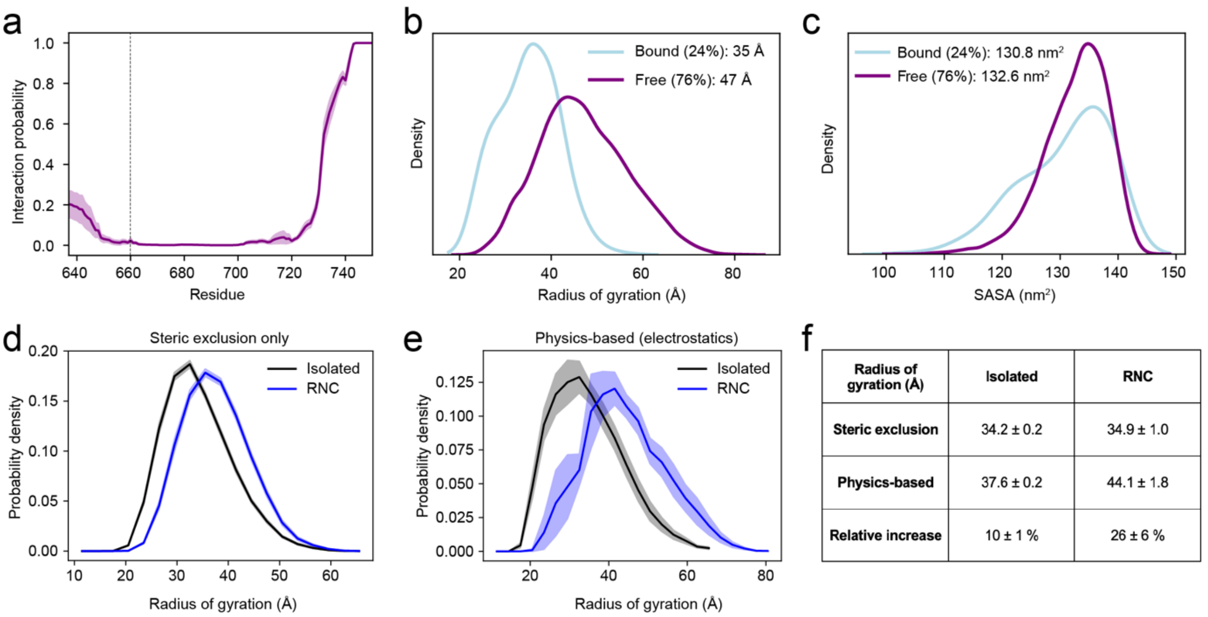


| Radius of gyration (Å) | Isolated | RNC |
|---|---|---|
| Steric exclusion | 34.2 ± 0.2 | 34.9 ± 1.0 |
| Physics-based | 37.6 ± 0.2 | 44.1 ± 1.8 |
| Relative increase | 10 ± 1 % | 26 ± 6 % |

**Figure S6.** Ribosome interactions and compaction of the unfolded state on the ribosome. **a.** Ribosome-nascent chain interaction probability of an unfolded FLN5 RNC (FLN5+31) plotted along the protein sequence (mean ± s.e.m.) from previous work[7]. **b.** Distributions of radius of gyration of unfolded FLN5 (FLN5+31 RNC) for the subensembles with the N-terminus (residues 637-660) bound to the ribosome and free. **c.** Distributions of the solvent-accessible surface area (SASA) of unfolded FLN5 (FLN5+31 RNC) for the subensembles with the N-terminus (residues 637-660) bound to the ribosome and free. **d-e.** Probability distributions (mean ± s.e.m.) of unfolded FLN5 on and off the ribosome obtained with an all-atom, steric exclusion-only model (d) and physics-based force field integrated with experimental NMR data (e) from previous work[7]. **f.** Ensemble-averaged radii of gyration of unfolded FLN5 and relative increase on the ribosome compared to isolated FLN5 (mean ± s.e.m.). Both isolated ensembles (steric- and physics-based models) are in excellent agreement with the radius of gyration derived from SAXS experiments (34.4. ± 0.6 Å)[7]. The significantly larger structural expansion (increase in radius of gyration) on the ribosome obtained with the physics-based approach shows that the expansion is driven by electrostatic repulsion between the negatively charged ribosome surface and unfolded state, given that the isolated ensembles have an identical radius. Van der Waals interactions between NC and ribosome atoms would be expected to lead to compaction on the ribosome as these interactions are attractive. Likewise, $Mg^{2+}$ ions that surround the ribosome surface would be expected to result in compaction by attracting the negatively charged unfolded state, which is not observed. This agrees with $^{19}F$ NMR experiments that previously demonstrated that $Mg^{2+}$ ions have negligible effects on NMR linewidths and folding thermodynamics of FLN5[13].

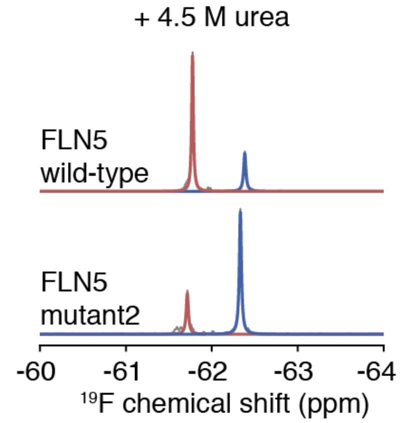


**Figure S7.** $^{19}$F NMR spectra of FLN5 655tfmF (wild-type and mutant 2) in 4.5 M urea recorded at 298 K and 500 MHz. Observed and fitted spectra shown in grey and colour, respectively.

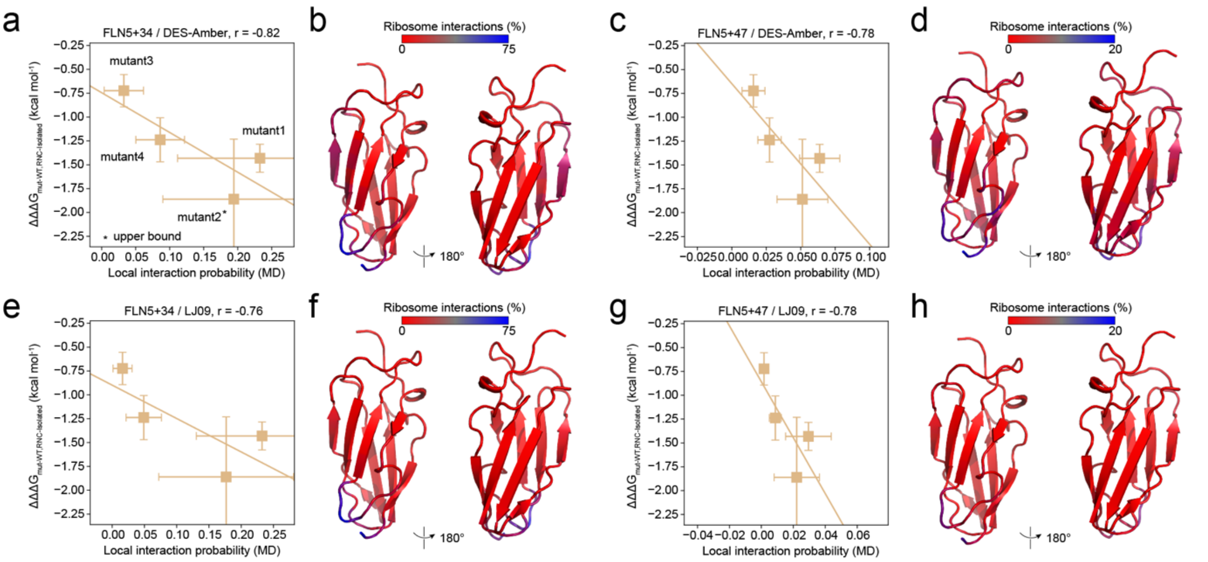


**Figure S8.** Correlation between ribosome-induced mutant (de)stabilisation and local ribosome interaction probabilities as in Fig. 3e-f. Correlations were plotted and local interaction probabilities mapped for FLN5+34 / DES-Amber (panels a-b), FLN5+47 / DES-Amber (panels c-d), FLN5+34 / LJ09 (panels e-f), and FLN5+47 / LJ09 (panels g-h).

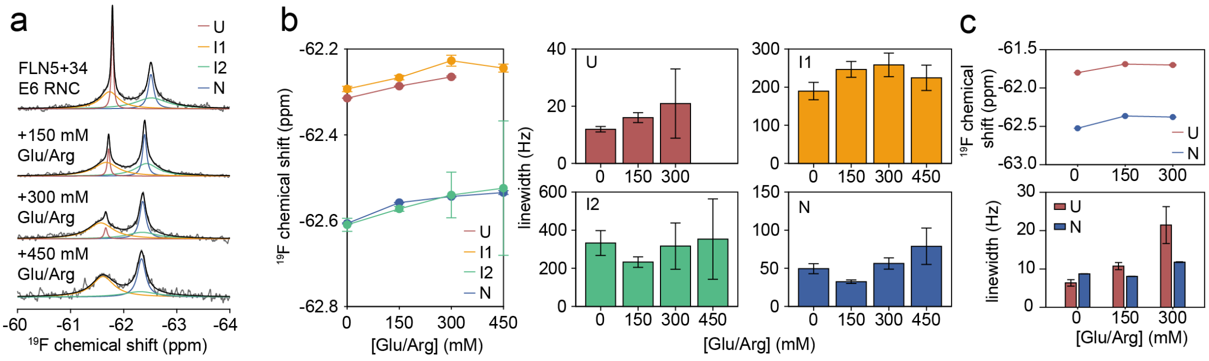


**Figure S9.** Salt titration of FLN5 E6 on and off the ribosome. **a.** $^{19}$F NMR spectra of FLN5+34 E6 in increasing concentrations of L-Glu/L-Arg recorded at 298 K and 500 MHz. Observed, fitted and total fitted spectra shown in grey, colour and black, respectively. **b.** $^{19}$F chemical shift and linewidth changes of all FLN5+34 E6 RNC conformational states with increasing ionic strength (mean ± s.e.m.). **c.** $^{19}$F chemical shift and linewidth changes of isolated FLN5Δ2 E6 conformational states with increasing ionic strength (mean ± s.e.m.).

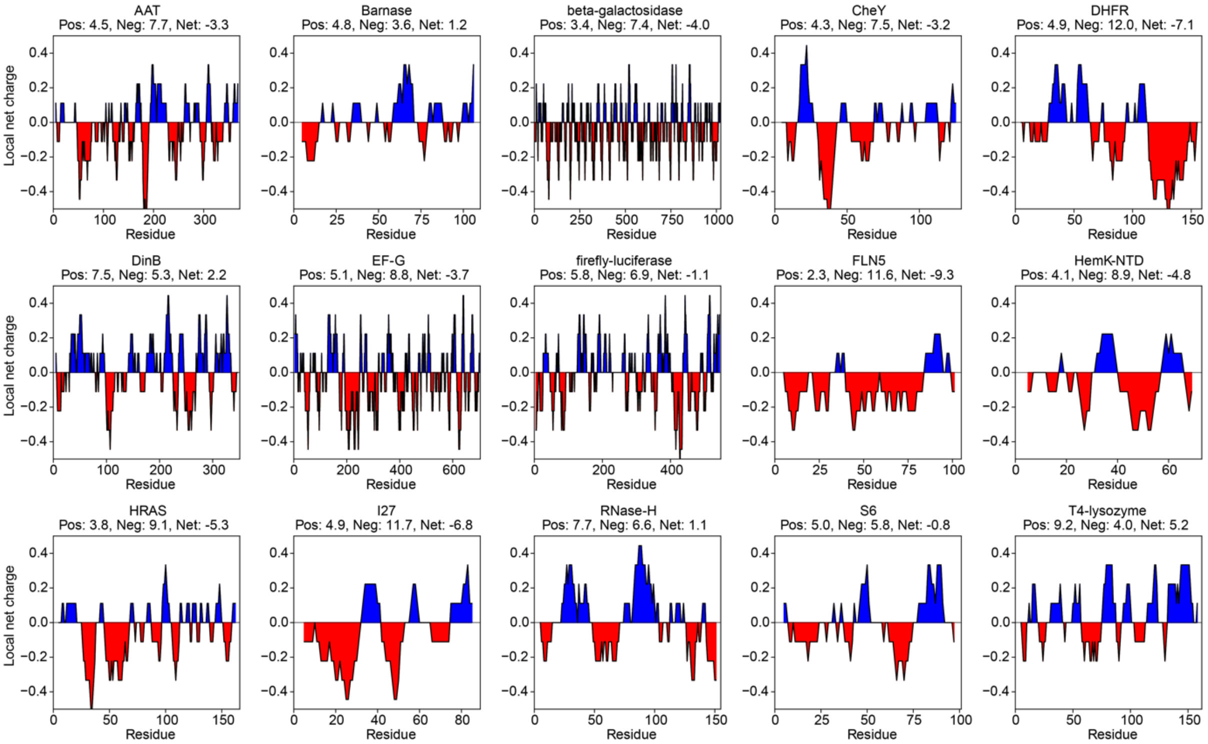


**Figure S10.** Local net charge of proteins studied on the ribosome in the literature (Table S2) averaged over a window size of nine residues. The surface positive (Pos) and negative (Neg) surface areas are annotated above each plot and their difference (Net). Charge distribution throughout the sequence appears to relate to decreased folding rates in T4 lysozyme and EF-G[9,51,103], mediated by ribosome interactions with positively charged patches in unfolded conformations as observed directly for T4 lysozyme and FLN5[9,18].

### Supplementary tables

| RMSD (Å) | All-atom | Backbone heavy atoms |
|---|---|---|
| **RNC vs. isolated** | 2.0 ± 0.3 | 1.0 ± 0.2 |
| **RNC vs. RNC** | 2.0 ± 0.3 | 1.0 ± 0.2 |
| **isolated vs. isolated** | 2.0 ± 0.3 | 1.0 ± 0.2 |

**Table S1.** Average (± standard deviation) of the pairwise RMSD matrix comparing structures on and off the ribosome, and within the ensembles separately. These were calculated using 10,000 structures for each ensemble.

| Protein/domain | Net charge at pH 7.4 | Native state stability / folding onset | Protein / domain length (residues) | Refs. |
|---|---|---|---|---|
| AAT | -10 | Destabilised | 372 | [27] |
| Barnase** | +3 | Destabilised | 110 | [19] |
| β-galactosidase | -40 | Destabilised; high population of coTF intermediates | 1,024 | [16] |
| Chemotaxis protein Y (CheY) | -4 | Destabilised; locally stabilised in a positively charged helical region (around residue Met 17) | 129 | [20] |
| DHFR**/*** | -10 | Destabilised; high population of coTF intermediates | 159 | [17,19,20] |
| DNA polymerase IV (DinB) | +9 | Native state globally unaffected; locally destabilised in a negatively charged region (around residue Met 249) | 351 | [20] |
| EF-G | -24 | Destabilised | 704 | [104] |
| Firefly luciferase | -6 | Destabilised; high population of coTF intermediates | 548 | [105] |
| FLN5 | -9 | Destabilised; high population of coTF intermediates | 105 | [7,13,18] |
| HemK NTD | -4 | Destabilised; high population of coTF intermediates | 73 | [14,15,106] |
| HRAS (G-domain) | -8 | Destabilised; high population of coTF intermediates | 166 | [7] |
| I27 | -6 | Destabilised; high population of coTF intermediates | 89 | [7] |
| RNase H** | +1 | Destabilised | 155 | [19,42] |
| S6 | 0 | Positively charged mutants are stabilised on the ribosome (earlier coTF onset). | 101 | [26] |
| T4 lysozyme* | +8 | Native state globally unaffected; intermediate states unaffected; reduced folding rate and unfolded state compacted due to ribosome interactions/electrostatic attraction. | 162 | [9] |

**Table S2.** Summary of stability measurements of RNCs in the literature. Stability refers to global stability of the native state relative to in isolation ($\Delta G_{t(N)}$) unless stated otherwise, inferred from unfolding kinetics or folding free energies ($\Delta G_{f(N),RNC}$). Net charge was calculated by assuming all Asp/Glu residues carry a charge of -1, Arg/Lys +1, and His 0.

*Unfolding transitions are indistinguishable on and off the ribosome but folding rates are slower on the ribosome. This can be modulated by distance from PTC and ionic strength (both increase folding rate).

**Ref. [19] shows that their destabilisation correlates with net charge: more negatively charged proteins are more destabilised. For example, barnase (+3) is only destabilised by +0.4 ± 0.1 kcal mol$^{1}$. These stability effects were observed to be dependent on the distance from the ribosome surface. Additionally, ref. [20] also showed that increasing distance from PTC and higher ionic strength decrease the destabilisation.

***Ref. [20] shows by means of ligand binding to DHFR that the ribosome can destabilise the native state without significantly altering its structure, in line with this work.
